# Multicomplex Integrative Structural Modeling of a Human Histone Deacetylase Interactome

**DOI:** 10.64898/2026.03.16.712198

**Authors:** Jules Nde, Kartik Majila, Rosalyn C. Zimmermann, Cassandra Kempf, Ying Zhang, Joseph Cesare, Janet L. Thornton, Jerry L. Workman, Laurence Florens, Shruthi Viswanath, Michael P. Washburn

## Abstract

Histone Deacetylase (HDAC) 1 and 2 are key enzymatic components in multiple large chromatin remodeling complexes including NuRD, SIN3, and CoREST. In addition, both HDAC 1 and 2 contain a large intrinsically disordered region (IDR) within their C-terminal domain (CTD). How HDAC1/2 assemble into these complexes and the structure of the CTD IDR remains poorly understood. Here, we used HDAC1/2 to isolate their protein interaction networks from cells and used crosslinking mass spectrometry (XL-MS) coupled with the Integrative Modeling Platform to build structural models of the NuRD, SIN3A, and CoREST complexes. Next, we implemented an AlphaFold-enabled XL-MS constrained modeling approach to investigate how HDAC1 could assemble into these complexes. We show that the CTD IDR of HDAC1 folds into alpha helices in these complexes. Finally, we built a complete integrative structural model of a NuRD subcomplex including the abundant HDAC1:MBD3:MTA1:GATAD2B:RBBP4 subunits, which included 6 IDRs. The approaches used herein are broadly applicable for the study of protein complexes and protein interaction networks that can provide important insights into IDRs.

## Introduction

The class I histone deacetylases (HDAC) HDAC1 and 2 are primarily localized to the nucleus and have strong deacetylase activity ^1^. Linked to many diseases ^2^, including nearly all cancers ^3^, HDAC1/2 carry out their functions as part of large transcriptional corepressor complexes including CoREST, MIER, NuRD, and SIN3^4^. HDAC1/2 regulate a variety of cellular processes such as cellular and DNA metabolism, cellular differentiation, DNA damage response, cell cycle regulation, proliferation, and cellular differentiation ^1^. Based on their high 85% sequence similarity, HDAC1 and HDAC2 are often assumed to function in an identical manner. However, studies have identified distinct functions between the paralogues where they demonstrate distinct roles in regulating and promoting differing stages of development, stem cell pluripotency, and cancer progression ^5–7^.

Given their importance in normal and diseased states, both proteins have had large portions of their structure crystallized, from amino acids (AAs) 8-376 and 1-367, for HDAC1 and 2, respectively ^8,9^. The crystallization consists primarily of the HDAC domain (AAs 8-376 and AAs 9-378), which includes the active sites at H140/141 or H141/142, for HDAC1 and 2, respectively. However, the last *ca.* 100 amino acids defining the C-terminal domain (CTD) of each protein with the highest regions of sequence dissimilarity ^10^ remain unresolved, within HDAC1 and HDAC2 Xray structures ending at Ala 376 ^11–13^ and Pro 375 ^14,15^, respectively. As a result, 22% of HDAC1 and 23% of HDAC2 remain structurally uncharacterized. The CTDs of HDAC1 (AA 390-482) and HDAC2 (389-488) are considered intrinsically disordered ^10^. Intrinsically disordered regions (IDR) play key roles in biological systems ^16–18^ but are challenging to study using current structural biology approaches^16,19^.

In addition, multiple subunits of the HDAC1/2-containing complexes also have IDRs ^10^. For example, recent studies of NuRD subunits have focused on how the CHD family members ^20^ and MBD2 interact with NuRD core ^21^. Other key proteins in the NuRD complex include the methyl-CpG-binding domain protein MBD3 ^22^, metastasis-associated protein MTA1 ^23^, GATA zinc finger domain containing 2B GATAD2B ^24^, and the retinoblastoma binding protein RBBP4^25^. This core module forms the enzymatic and scaffolding foundation onto which other subunits may be incorporated ^12,26,27^. However, how the IDRs within NuRD fold into complete structures remains poorly understood.

Integrative structural modeling (ISM) is a powerful approach to study protein complexes and a rapidly growing field ^28,29^. We recently described a fully resolved integrated structural model of the *S. cerevisiae* Sin3 large (Sin3L) complex that combined cryo-EM and cross-linking mass spectrometry^30^. In this study, native Sin3L complexes were affinity purified from *S. cerevisiae* and high resolution cryo-EM maps of native Sin3L complexes were obtained^30^. Integration of XL-MS and cryo-EM resulted in a fully resolved Sin3L structural model containing 12 subunits that expanded the overall amino acid coverage from previous models from ∼50% to 100%^30^. An important contribution to this expansion of sequence coverage was the ability of XL-MS to detect and identify cross-links in IDRs and model IDRs, including those in the *S. cerevisiae* Sin3L complex components of ASH1, CTI6, PHO23, SAP30, SDS3, and RPD3. RPD3 is the *S. cerevisiae* ortholog of HDAC1 and HDAC2, and RPD3 has a C-terminal domain IDR from amino acids 388-433^30^. In the reported structure of *S. cerevisiae* Sin3L we in fact reproducibly detected and identified cross links between lysine 415 in RPD3 and different lysines -in RPD3, PHO23, RXT3, and SIN3^30^. These results on the *S. cerevisiae* Sin3L complex demonstrate the ability of XL-MS to detect and identify IDRS, in addition to demonstrating our ability to build integrative structural models of IDRs in proteins.

We have also used affinity purifications and crosslinking mass spectrometry (XL-MS) coupled with ISM to study a human SIN3A subcomplex ^31^, the human SPIN1:SPINDOC complex ^32^, and the human WDR76:SPIN1 complex ^33^. Here, we applied these approaches to build structural models of HDAC1/2 containing complexes from endogenous HDAC1/2 protein complexes. Affinity purifications of HDAC1 and HDAC2 were analyzed using XL-MS yielding extensive and specific sites of protein-protein interactions in each system. We then used the Integrative Modeling Platform ^34^, which has been used to study NuRD subcomplexes ^35^, to build structural models of portions of the HDAC1 protein interaction networks including NuRD ^4,35,36^, SIN3A^4,37^, and CoREST^4,38,39^. Next, we used AlphaFold3 ^40^ and XL-MS constrained HADDOCK docking ^41^ to determine how HDAC1, including the previously unresolved C-terminal IDR, behaves in this system. Lastly, using the crosslinks we were able to build a complete model of a NuRD subcomplex containing HDAC1:MBD3:MTA1:GATAD2B:RBBP4 that contains a total of six IDRs.

## Results

### XL-MS Based Protein Interaction Networks of HDAC1 and HDAC2

HDAC1 and HDAC2 share the most sequence similarity of the four Class I HDACs (Fig. S1A). To gain insight into each protein interaction and structure, we performed XL-MS on C-terminally Halo-tagged HDAC1 and HDAC2 stably expressed in HEK293T cells. Previous work from our lab has shown that the C-terminally tagged HDACs maintain nuclear localization and interaction with known binding partners, as opposed to an N-terminal tag which limited interactions to chaperones and prefoldin complex members ^4^. The six XL-MS datasets were searched independently using the XlinkX node in Proteome Discoverer 2.4. Combining the three HDAC1-Halo XL-MS datasets, a total of 1471 crosslinks spectrum matches (CSMs) passed the 1% FDR cut-off (Table S1A), 614 for intra-molecular crosslinks (intra-XLs) and 857 inter-molecular crosslinks (inter-XLs). Slightly less CSMs were detected combining the three HDAC2-Halo XL-MS datasets adding up to 1187 total CSMs breaking into 666 and 521 intra-and inter-XLS, respectively (Table S1B). For HDAC1, 17 peptides carried crosslinked lysine residues, covering 55% of its sequence, while 13 HDAC2 peptides were detected with crosslinked lysine residues, mapped to 40% of its sequence (Fig. S1B). Twelve sites were detected on lysines conserved in HDAC1 and HDAC2; while the peptide bearing HDAC2-K166 was not detected as crosslinked in the HDAC2 datasets and HDAC1-K126 aligns with an arginine residue in HDAC2, which cannot be crosslinked by DSSO. Two non-overlapping peptides were detected within the less homologous CTDs, bearing HDAC1-K457 and HDAC2-K474 as crosslinked.

Four of the conserved lysines were located within peptides shared between HDAC1 and 2: K50/51, K89/90, K165/166, K200/201. When plotting the total CSMs identified for each of the crosslinked residues (Figure 1A), a limited number of CSMs (less than 5%) were mapped to HDAC2 peptides in the HDAC1-Halo pull-downs and vice versa. In other words, since each HDAC did not appear to pull-down a significant amount of the other isoform, we therefore considered the shared peptides detected in the HDAC1 XL-MS datasets to be likely HDAC1 peptides, and conversely. The only inter-molecular XLs recovered between HDAC1 and HDAC2 were HDAC1-K89_X_HDAC2-K75, in which K75 is within a peptide unique to HDAC2 in the HDAC1 dataset, and the reciprocal HDAC1-K74_X_HDAC2-K90 crosslink, in which K74 is within a peptide unique to HDAC1 in the HDAC2 dataset (Table S1C). These lysine residues are likely within the main site of homo and heterodimerization between HDACs, when it occurs. Overall, the intra-molecular crosslink profiles observed for HDAC1 and 2 were very similar (Fig. S1C), which was to be expected for proteins with high sequence and structural similarities.

**Figure 1:**
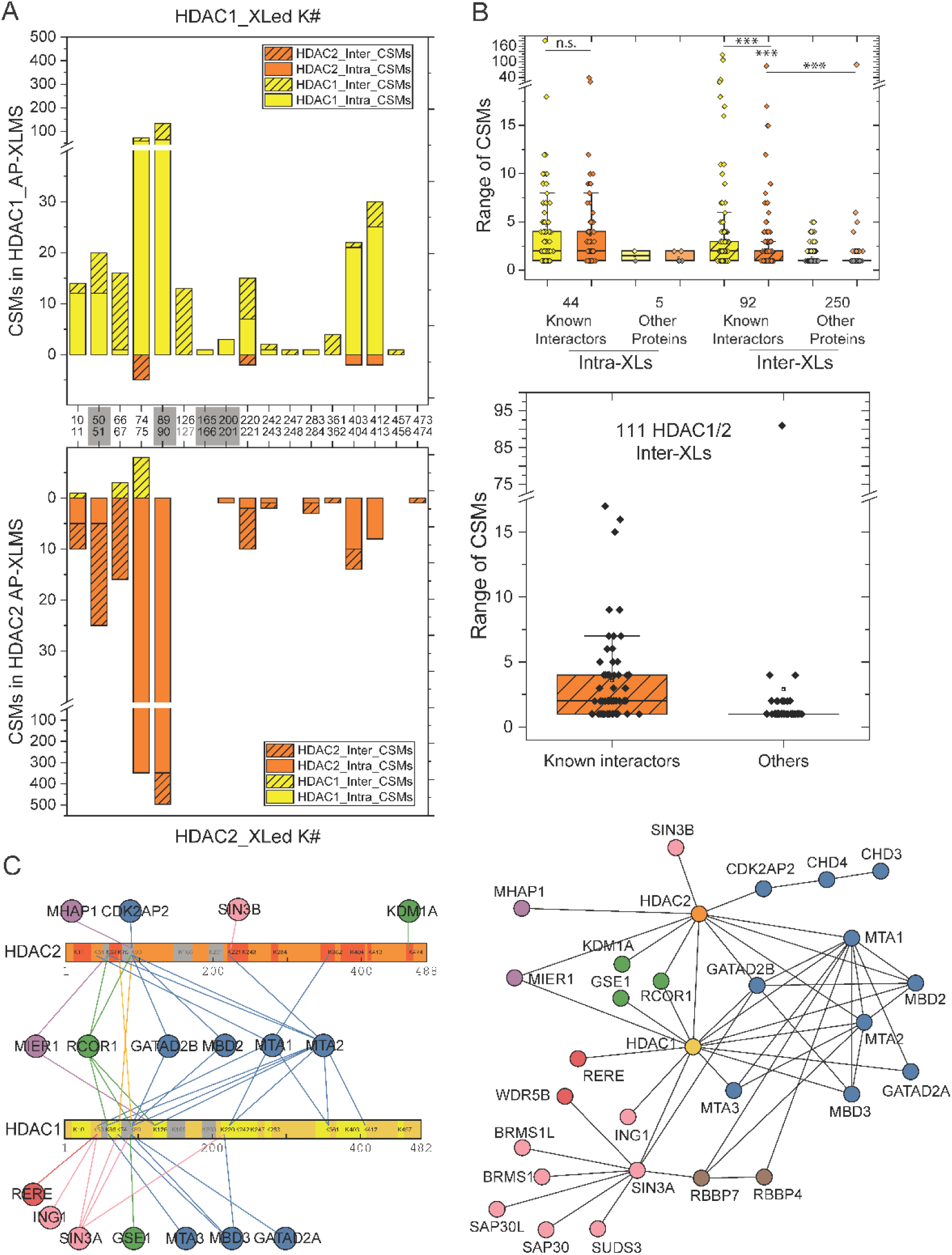
XL-MS defines HDAC1 and HDAC2 direct interaction networks. **A**. The total number of crosslink peptide matches (CSMs) mapped to HDAC1 and HDAC2 peptides in the HDAC1 and HDAC2 AP-XL-MS analyses are reported in the top and bottom panels, respectively (see Table S1C). HDAC1 and HDAC2 CSMs are reported in yellow and orange, while intra- and inter-crosslinks are plotted in solid and hashed bars. Lysine residues within peptides shared between the 2 isoforms are marked in grey along the x-axis (see Figure S1B and Table S1E). **B.** The upper plot reports the range of CSMs values measured for intra- and inter-XLs for known interactors of the HDAC1/2-containing complexes and/or nucleosomes and for proteins not previously reported as such (“Other”). Significant differences in CSM distributions were assessed using a Mann-Whitney test with *** denoting p-values <0.0001 (see Table S1E). The distribution of CSMs values for intermolecular crosslinks between HDAC1 and HDAC2 and their known interactors or other proteins are plotted in the lower panel. **C.** The intramolecular interactions between HDAC1/2 and their direct partners are plotted with xiView (left panel), while all interlinks between subunits of HDAC1/2-containing complexes are reported in the right panel (see Table S1E for input values).

Combining both HDAC1 and HDAC2 AP-XL-MS datasets resulted in 596 non-redundant crosslinks (Table S1C) reporting interactions between 359 distinct proteins (Table S1D). Crosslinked proteins were grouped based on their membership to known HDAC1/2-containing complexes (51 subunits), their previously reported interaction with HDAC1 and/or HDAC2 (13 “HDAC1/2-interacting” proteins) or with at least one of the 4 canonical histones (“HIST-interacting”). This group of 104 proteins represent known or likely interactors based on the IntAct ^42^, Corum ^43^, and Complex Portal ^44^ databases. Proteins not previously reported to interact with HDACs or histones were split into 2 groups: 64 were annotated as localizing to the nucleus (“NUC”), while the “OTHER” group contains the remaining 191 proteins. Not surprisingly, the range of CSMs observed between known interactors was significantly larger than the range of CSMs for other proteins, for both intra and inter-molecular crosslinks (Figure 1B, upper panel). While about half of the 111 unique inter-XLs involving either HDACs were with proteins not previously known to interact with either one, the range of CSMs for these XLs was lower than the CSMs measured between HDAC1/2 and known interactors (Figure 1B, lower panel), with a few outliers. This indicates that the Halo affinity purifications followed by DSSO crosslinking enriched for biologically relevant interactions.

Analyzing the XL-MS interactomes allowed us to identify HDAC1 and HDAC2 direct vs indirect interactors, a classification which was previously limited in our HDAC AP-MS studies ^4^. To visualize the large protein interactomes of both HDAC1 and HDAC2, we used the xiView platform ^45^. HDAC1 interacted directly with 13 members of 5 different HDAC1/2-containing complexes, while HDAC2 interacted with 10. Six of these subunits interacted with both HDACs, mostly at the same locations defining K50/51, K66/67, K74/75, and K89/90 as interactions hot spots (Figure 1C). In all 27 subunits of the COREST, NuRD, SIN3, MIER, and WHERE^46^ complexes had primary or secondary interactions with HDAC1 or 2 (Figure 1C). The analysis of HDAC1 and HDAC2 crosslinks identified several protein nodes (which are defined as proteins with ≥ 5 unique protein interactors) (Figure 1C). As expected, HDAC1 and HDAC2 exist as protein nodes. In addition, we observed several NuRD complex members (CHD4, GATAD2B, MBD3, MTA1, MTA2, and RBBP7), and MIER1, KDM1A, and SIN3A function as protein nodes in this interactome. On the one hand, these results allowed us to identify with greater detail how HDAC1 and 2 associate with proteins within their complexes. On the other hand, while many intermolecular crosslinks were observed within the nucleosome and between histones and both known and unknown interaction partners (Table S1C), no intermolecular crosslinks were detected between the canonical histones and any of the subunits of the HDAC1/2-containing complexes, including the baits HDAC1 and 2. This could indicate that the substrate of these chromatin remodeling complexes, the N-terminal histone tails, are difficult to crosslink using a crosslinker targeting primary amines, considering that most lysine side chains within the histone tails are modified.

Several of the HDAC complexes (CoREST, NuRD, and SIN3) have previous structural information for at least one complex member ^12,31,35–37,47–53^. Since XL-MS data is most powerful when combined with complementary structural information derived from multiple techniques, we next used Bayesian integrative structure determination *via* the Integrative Modeling Platform (IMP) ^34^. This allowed us to combine structural and biochemical data at several scales coupled with statistical analysis to determine the integrative structures of these complexes ^34,35^.

### Crosslink-guided assembly of HDAC1/2 within the NuRD complex

The NuRD complex had the most crosslinks compared to all other HDAC1/2-containing complexes we recovered in this study (Table S1). As a result, we had a greater number of crosslinks to build a model of the NuRD complex. The NuRD complex is a transcriptional regulator that can function as a corepressor or coactivator that functions by deacetylating histone tails and by ATP-dependent chromatin remodeling ^26^. HDAC1 and HDAC2 function as one of the enzymatic components of the complex, in which they bind to the ELM2-SANT domain within an MTA (MTA1/2/3) paralog (Fig. S2A) ^26^. The complex also contains the CHD3/4, MBD2/3, and GATAD2A/B paralogs and the CDK2AP1 subunit. Since they had the most crosslinks (Figure 2A), we used the HDAC1, GATAD2B, MBD3, MTA1, and RBBP4 paralogs as core members of the complex in our integrative structural modeling. Previous integrative structures of NuRD have been determined with several types of structural information including cryo-EM and negative-stain EM maps as well as X-ray and NMR structures, homology models, and XL-MS studies (Fig. S2C) ^35^. Our current study adds to the previous structural information by including direct intra- and inter-molecular crosslinks between the 5 core subunits (Fig. S2D), with a stoichiometry of 2 MTA1, 2 HDAC1, 4 RBBP4, 1 MBD3, and 1 GATAD2B subunits (Fig. S2B), which provides further clarity to the histone deacetylase module of the NuRD complex.

**Figure 2:**
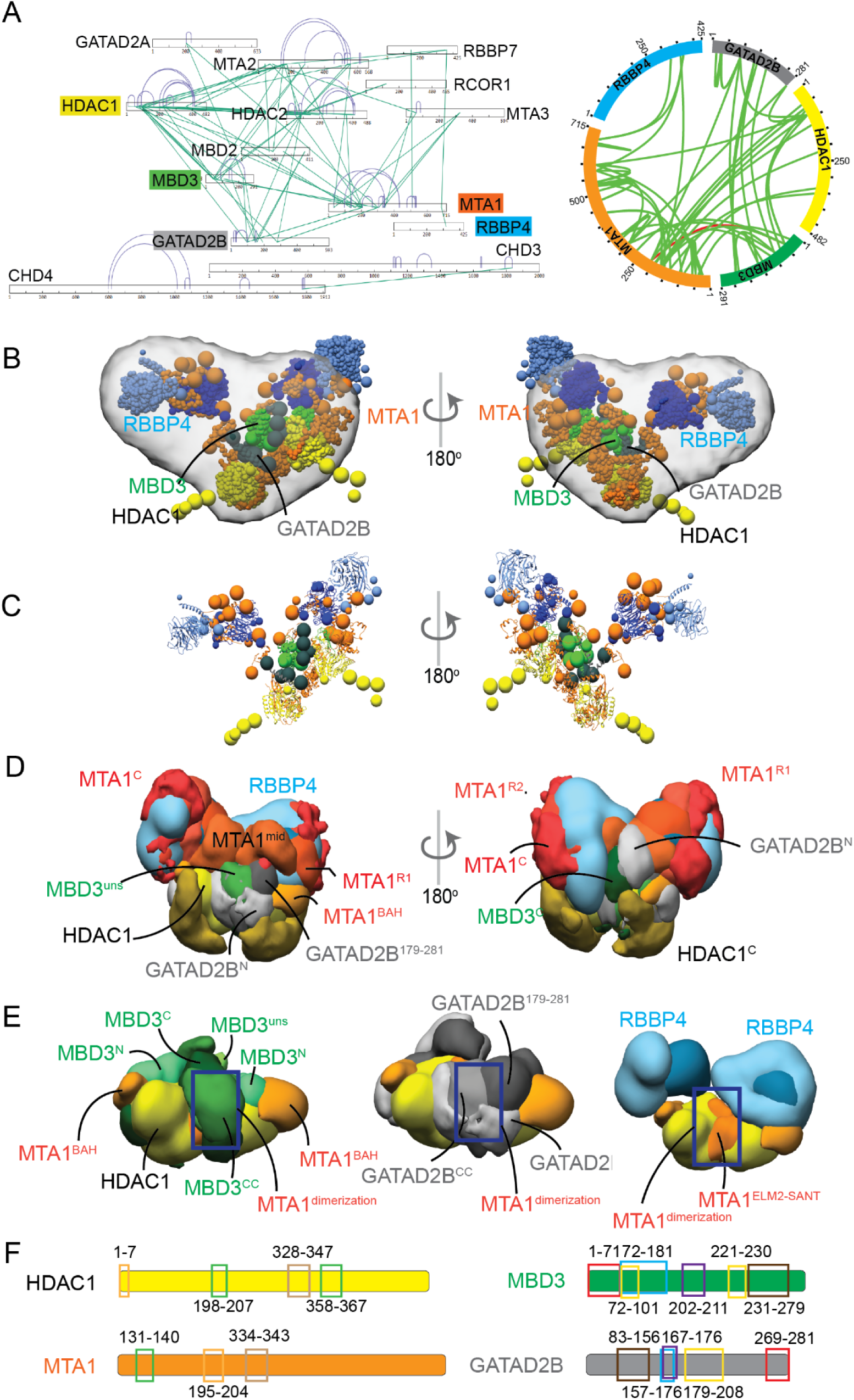
Integrative modeling of the NuRD complex. **A.** xiView visualizations of the crosslinks observed for members of the NuRD complex. The linear diagram reports all observed crosslinks (intra- and inter-molecular in purple and green, respectively). The Circos plot reports the crosslinks between the 5 subunits used for integrative modeling, with links that are satisfied in the ensemble of models from the major cluster in green and unsatisfied in red. **B.** Representative bead model from the most populated cluster of integrative models for the NuRD complex, shown with the NuRD EM map. The model is colored by subunit. **C.** Representative bead model from panel B with regions of known structure shown in ribbon representation. **D.** Localization probability density maps showing the position of different domains/subunits in the cluster. The map specifies the probability of any volume element being occupied by a domain in the ensemble of superposed models from the cluster. The domain densities are colored according to Figure S2A. **E.** Localization probability density maps showing the MTA1-HDAC1 dimer with MBD3 (left), GATA2B (center), and RBBP4 (right). Other protein domains are hidden for clarity. All maps are contoured at ∼10-20% (30% for MTA1^mid^, HDAC1^C^) of their respective maximum voxel values. **F.** Novel protein-protein interactions identified from contact maps of the major cluster (interacting domains are represented in the same color). See Table S2 for the complete list of interaction interfaces within the modeled NURD complex and Figures S2-S3 for details about the structural information used by IMP, sampling exhaustiveness protocol, and precision analysis.

Integrative modeling of the NuRD complex produced effectively a single cluster of models (97% of a total of 28,914 models) with a model precision of 55 Å, in which we define model precision as the average RMSD between the cluster centroid and models in the cluster (Fig. S3A-F). We find that 99% of the input crosslinks are satisfied (Figure 2A-right). The cross-correlation of the localization density map for the models in the major cluster with the EM map was also high at 0.86 (Figure 2B). Furthermore, the similarity of our current NuRD structure to a previously determined structure using an independent set of crosslinks validated our approach ^35^. Within the complex, the MTA1-HDAC1 dimer forms the scaffold for the complex, fitting obliquely in the EM map, with the MTA bromo-adjacent homology (MTA1^BAH^) domains flanking it on either side (Figure 2B-E). MTA1^mid^, which connects the N-terminal and C-terminal regions of MTA1, is located near the MTA1 dimerization interface (Figure 2D). Its localization is poor, *i.e.* the localization density is spread out, indicating it could be flexible and heterogenous. The C-terminal half of each MTA1 (MTA1^467–715^ containing MTA1^R1^, MTA1^R2^, MTA1^C^) extends like two arms from the MTA1-HDAC1 dimer and is similarly heterogenous (Figure 2D). HDAC1^C^ is located at the base of the MTA1-HDAC1 dimer, and it is also localized poorly (Figure 2D). The N-terminal MBD3 methyl-CpG-binding domain (MBD3^N^) is localized at two symmetric binding sites on the MTA1-HDAC1 dimer and is buried beneath the C-terminal RBBP binding regions of MTA1 and GATAD2B^179-281^ (Figure 2D-E). The unstructured region of MBD3 (MBD3^uns^), which contains an IDR, is at the MTA1 dimerization interface (Figure 2D-E). The coiled-coil domains of the MBD3-GATAD2B (MBD3^CC^-GATAD2B^CC^) complex are located on top of the MBD3^uns^ at the MTA1 dimerization interface (Figure 2D-E). MBD3^C^ winds up on top of and around the MTA1 dimerization interface, though it is localized poorly (Figure 2D-E). The GATAD2B domains are packed on top of MBD3 domains and spread across the MTA1-HDAC1 dimer (Figure 2E). GATAD2B^N^ is near the MTA1-HDAC1 dimerization interface, while GATAD2B^179-281^ is on top of the MBD3^MBD^ domain on the MTA1-HDAC1 dimer (Figure 2D-E). These localizations are consistent with the predicted localization of GATAD2 in NuRD ^35,36^.

Precision analysis of the NuRD integrative structure shows that the MTA1-HDAC1 dimer, MTA1^BAH^ and one of the MTA1-RBBP4 dimers are localized at high precision, while MBD3, GATAD2B, MTA1^mid^, and HDAC1^C^ are localized with low precision (Fig. S3G). Additionally, we have summarized a list of newly predicted protein-protein contact regions that include regions not covered by existing structures (Figure 2F, Table S2). Several domains, such as MTA1^BAH^, MTA1^mid^, MTA1-RBBP4 complexes, HDAC1^C^, GATAD2B^N^, and GATAD2B^179-281^ are exposed, indicating they could potentially interact with nucleosomal DNA or other proteins. Overall, the positions of these domains are consistent with their positions in a previously determined structure ^35^.

### Crosslink-guided assembly of HDAC1/2 within the SIN3 complex

The SIN3 complex functions as a corepressor, utilizing the deacetylation activity of HDAC1/2 to repress transcription of specific genes ^37^. The core scaffolding components, SIN3A and SIN3B, play roles in regulating cell cycle and tissue development ^37^. The two paralogues have differing roles in regulating cancer progression as SIN3A loss is associated with an increase in metastasis, while SIN3B loss appears to decrease metastatic capacity ^54,55^. In our XL-MS data, SIN3A had the most inter-complex interactions, while SIN3B was only crosslinked to HDAC2 (Figure 3A). Several of our XL-MS results are validated by the previously reported SIN3A/B XL-MS results ^37^, such as HDAC1/2-SIN3A interactions occurring at K50/51, K74, and K89/90 and HDAC2-SIN3B at K221 in both datasets. SIN3A and SIN3B interacted with HDAC2 at the same site (K221), suggesting that HDAC2 would have to be specifically involved in either a SIN3A or SIN3B complex.

**Figure 3:**
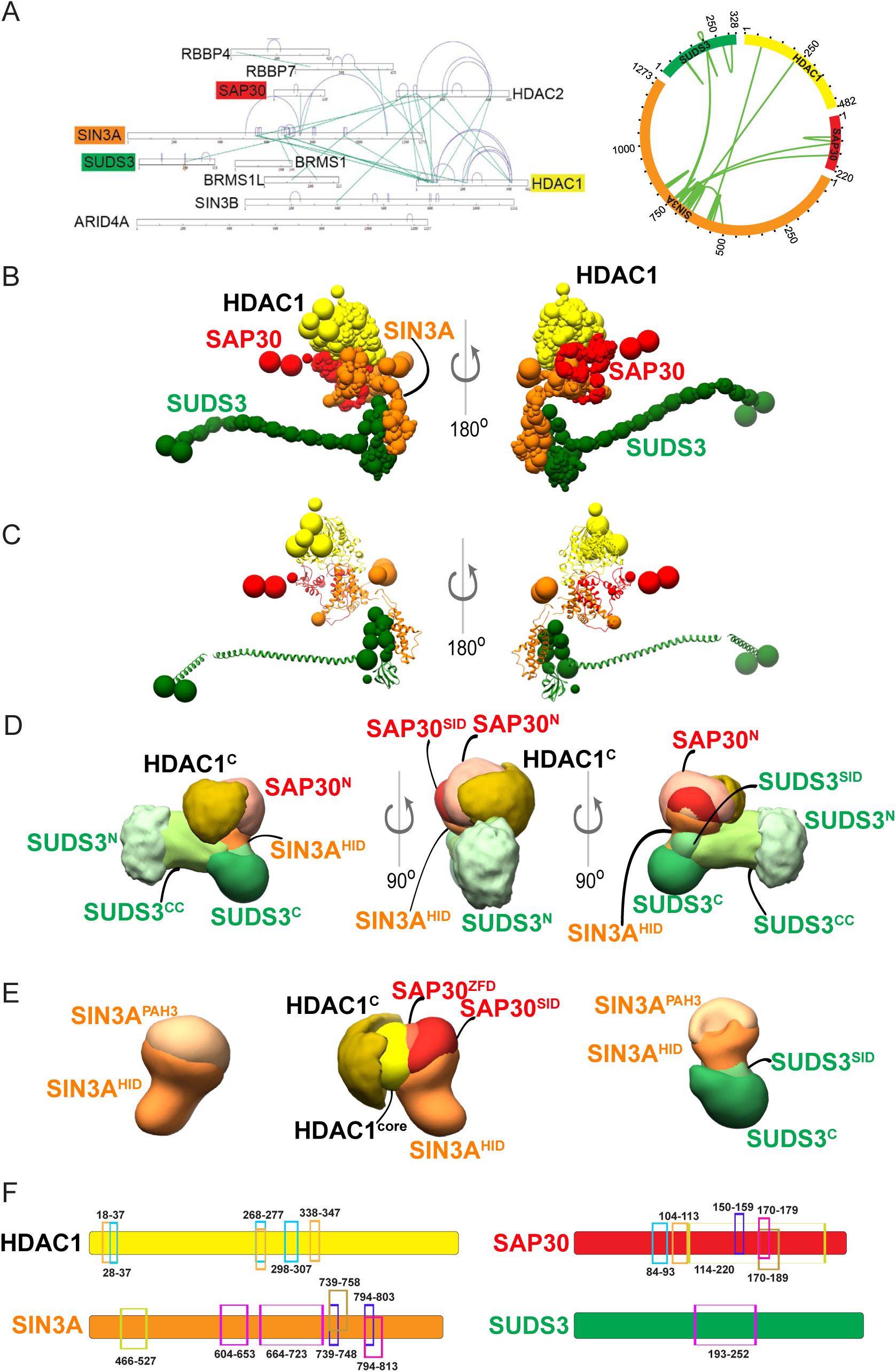
Integrative modeling of the SIN3A complex. **A.** xiView visualizations of the crosslinks observed for members of the SIN3A complex. The linear diagram reports all observed crosslinks (intra- and inter-molecular in purple and green, respectively). The Circos plot reports the crosslinks between the 4 subunits used for integrative modeling, with links that are satisfied in the ensemble of models from the major cluster in green and unsatisfied in red. **B.** Representative bead model from the most populated cluster of integrative models for the SIN3A complex. The model is colored by subunit. **C.** Representative bead model from panel B with regions of known structure shown in ribbon representation. **D.** Localization probability density maps showing the position of different domains/subunits in the cluster. The map specifies the probability of any volume element being occupied by a domain in the ensemble of superposed models from the cluster. **E-F.** Localization probability density maps of the core hub of SIN3A showing the scaffold of SIN3A^HID^ and SIN3^PAH3^ (E-left) along with HDAC1 and SAP30 domains (E-right) and SUDS3 domains (F) that are in the core hub. The domain densities are colored according to Figure S4A. These maps are contoured at ∼10% (except: SIN3A^HID^, SAP30^N^ are at 10%, and SUDS3^N^ and HDAC1^C^ are at 28%) of their respective maximum voxel values. **G.** Novel contacts identified at an average distance threshold of 10 Å between two beads across all the models in the major cluster (interacting domains are represented in the same color). See Table S2 for the complete list of interaction interfaces within the modeled SIN3A complex and Figures S4-S5 for details about the structural information used by IMP, sampling exhaustiveness protocol, and precision analysis.

Other members of the complex, ARID4A/B, BBX, BRMS1/L, FAM60A, FOXK1, ING1/2, RBBP4/7, SAP30/L, SAP130, and SUDS3^37^, were also detected in our XL-MS dataset with various numbers of intra and inter-molecular crosslinks (Figure 3A). Our integrative structural model then consists of a 4-subunit subcomplex of SIN3A in complex with HDAC1, SAP30, and SUDS3, built using the available crosslinks and structural information for each paralog (Fig. S4A-D). Integrative modeling of this SIN3A sub-complex produced effectively a single cluster (87% of a total of 29,602 models) with a model precision of 33 Å (Fig. S5A-F), in which 97% of the input crosslinks were satisfied (Figure 3A-right). The integrative structure indicates that SUDS3 spans the length of the complex from left to right, while the other three proteins are localized on one end (Figure 3B-D).

The core hub of the complex is at one end of SUDS3^CC^, with several protein domains located close to each other compactly, including the SIN3-interacting domain of SUDS3 (SUDS3^SID^), the paired amphipathic helix (SIN3A^PAH3^) domain, the SIN3A HDAC-interacting domain (SIN3A^HID^), the SAP30 zinc-finger domain (SAP30^ZFD^), and SIN3-interacting domain of SAP30 (SAP30^SID^) (Figure 3B-D). SIN3A^HID^ in the core hub forms a scaffold enveloping HDAC1 (Figure 3D-E). SAP30^ZFD^, HDAC1, and SUDS3^SID^ are mostly buried in the hub, while SAP30^SID^, co-located with SIN3A^PAH3^, is partly exposed (Figure 3D-E). Lastly, SUDS3^C^, SAP30^N^, and HDAC1^C^, extend from the hub (Figure 3D-E).

Precision analysis of the SIN3A monomer sub-complex indicates that HDAC1, parts of SIN3A^HID^, SAP30^ZFD^, SUDS3^CC^, and SUDS3^SID^ are localized at high precision, while HDAC1^C^, SAP30^N^, and SUDS3^N^ are localized at low precision (Fig. S5G). We also have summarized a list of newly predicted protein-protein contacts that include regions not covered by existing structures (Figure 3F, Table S2). Importantly, SIN3A forms several interactions with SUDS3 and SAP30. These interactions may explain how these proteins assist in stabilizing the SIN3A-HDAC1 interaction and help position HDAC1 correctly on the nucleosome. Since SIN3A does not have any DNA-binding motifs, it needs to be docked to DNA by other proteins in the complex, such as SAP30 and SUDS3. The domains of SUDS3 and SAP30 that are partly exposed, such as SUDS3^N^, SUDS3^C^, and SAP30^N^, may act as sites for binding transcription factors and/or DNA.

### Crosslink-guided assembly of HDAC1/2 within the CoREST complex

The CoREST complex acts as a transcriptional corepressor that plays important roles in cancer and neurodegenerative diseases^56^. The complex consists of three RCOR paralogs: RCOR1, RCOR2, and RCOR3. All three of the RCOR paralogs function as transcriptional repressors, with RCOR1 having the greatest repressive function ^57^. In addition to HDACs, KDM1A/LSD1, GSE1, HMG20A, HMG20B, PHF21A, ZNF217, ZMYM2, and ZMYM3 are also known members of the CoREST complex ^4^.

Our XL-MS data identified a network that consists of HDAC1 and HDAC2, RCOR1 and RCOR2, GSE1, and KDM1A (Figure 4A). Other members, HMG20A, RCOR3, and ZMYM3 had intra-protein XLs, while others were not detected in either HDAC1 or HDAC2 XL-MS experiments (HMG20B, PHF21A, ZNF217, and ZMYM2). It is known that KDM1A/LSD1 functions as a histone demethylase within the complex, in which it removes methyl groups on histone tails for transcriptional regulation ^58^. An 852 AA protein, KDM1A consists of a complex-stabilizing and nucleosome targeting SWIRM domain, two enzymatically active amine oxidase domains (AOD), and a protein-interaction TOWER domain (Fig. S6A) ^58,59^. RCOR1 consists of an ELM/SANT1 domain, known to mediate histone deacetylase recruitment, as well as a SANT2 domain that binds KDM1A and may function in binding the RCORs to nucleosomal DNA (Fig. S6A) ^60^. In our data, we find that HDAC1 and HDAC2 interact with RCOR1 at the same site (HDAC1/2 K89/90, respectively), which occurs within RCOR1 ELM2 binding domain (AAs 105-160). HDAC1 also binds RCOR2 within its ELM2 domain (AAs 46-100). These XL sites hence validate the known function of the ELM2 domain for HDAC recruitment to the scaffolding RCOR proteins ^57^.

**Figure 4:**
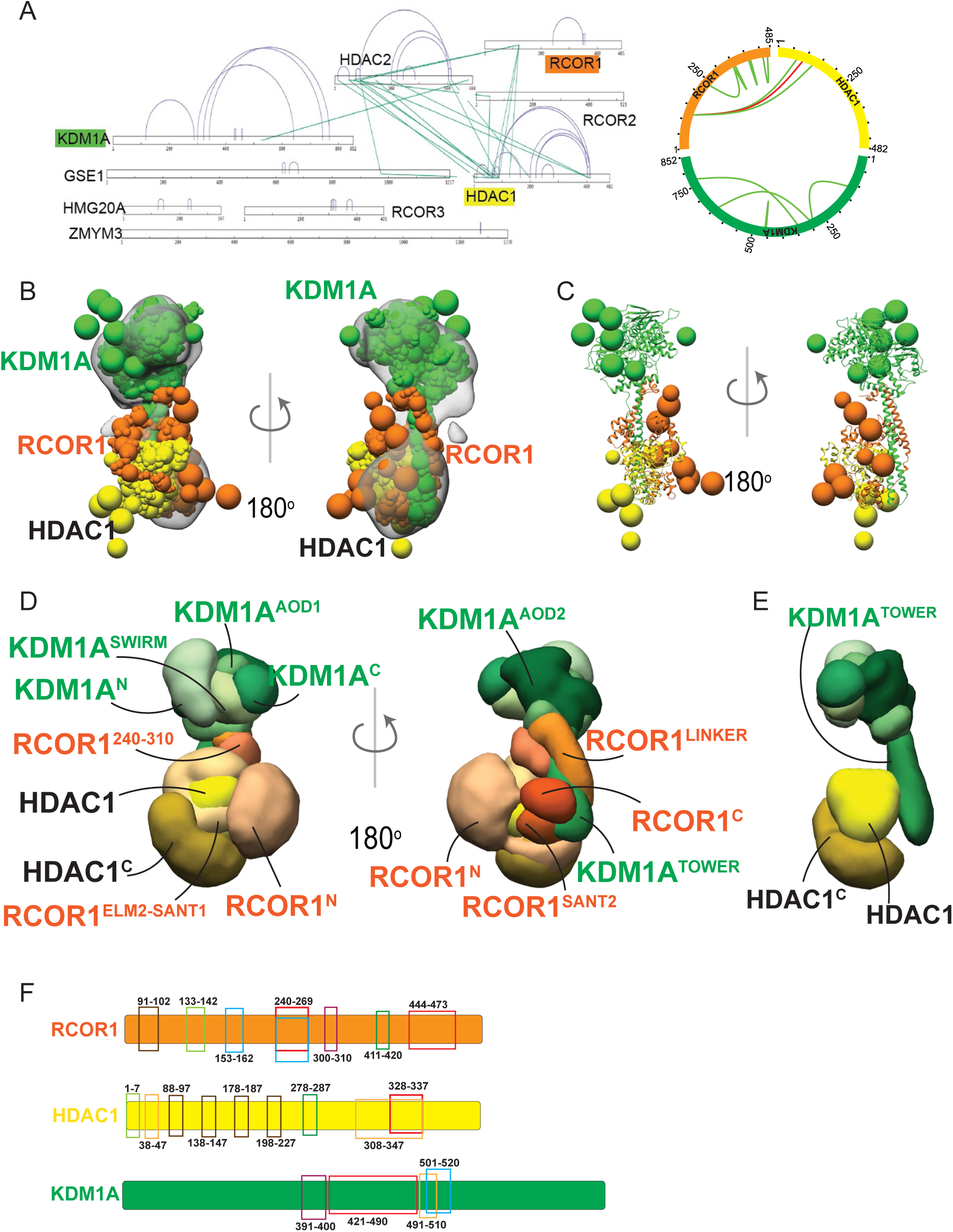
Integrative modeling of the CoREST complex. **A.** xiView visualizations of the crosslinks observed for members of the CoREST complex. The linear diagram reports all observed crosslinks (intra- and inter-molecular in purple and green, respectively). The Circos plot reports the crosslinks between the 3 subunits used for integrative modeling, with links that are satisfied in the ensemble of models from the major cluster in green and unsatisfied in red. **B.** Representative bead model from the most populated cluster of integrative models for the CoREST complex, shown with the CoREST EM map. The model is colored by subunit. **C.** Representative bead model from panel C with regions of known structure shown in ribbon representation. **D.** Localization of probability density maps showing the position of different domains/subunits in the cluster. The map specifies the probability of any volume element being occupied by a domain in the ensemble of superposed models from the cluster. The domain densities are colored according to Figure S6A. These maps are contoured at ∼10-15% of their respective maximum voxel values. **E.** Localization probability density maps for KDM1A-HDAC1. **F.** Novel contacts identified at an average distance threshold of 10 Å between two beads across all the models in the major cluster (interacting domains are represented in the same color). See Table S2 for the complete list of interaction interfaces within the modeled CoREST complex and Figures S6-S7 for details about the structural information used by IMP, sampling exhaustiveness protocol, and precision analysis.

We performed CoREST integrative modeling with HDAC1, KDM1A, and RCOR1 (Fig. S6A-D). Integrative modeling of the CoREST complex effectively produced a single cluster of models (69% of a total of 16,055 models) with a model precision of 12 Å (Fig. S7A-F), in which 92% of the input crosslinks were satisfied (Figure 4A-right). In addition, the cross-correlation of the localization density map to the EM map was 0.93. Precision analysis of the CoREST integrative structure demonstrates that the RCOR1-KDM1A complex and RCOR1-HDAC1 complex are localized at high precision while RCOR1^N^, HDAC1^C^, and KDM1A^N^ are at low precision (Fig. S7G).

Overall, this 3-subunit CoREST sub-complex forms a bi-lobed structure with most of KDM1A occupying the upper lobe, HDAC1 occupying the lower lobe, and the RCOR1-KDM1A complex connecting the two lobes (Figure 4B-D). The RCOR1-KDM1A complex comprises of the RCOR1 LINKER, and SANT2 domains interacting with KDM1A TOWER domain. KDM1A spans the length of the CoREST complex (Figure 4E). The RCOR1 N-terminus (RCOR1^N^) is localized at low precision towards the HDAC1-end of the structure (Figure 4B-C). The RCOR1^ELM2-SANT1^ region that forms a complex with HDAC1 also appears to interact with the KDM1A^TOWER^ domain (Figure 4D). RCOR1 AAs 240-310 (RCOR1^240-310^) are situated above the RCOR1^ELM2-SANT1^-HDAC1 complex and connects the RCOR1-HDAC1 complex with the RCOR1-KDM1A complex. RCOR1 also interacts with KDM1A AOD1 (KDM1A^AOD1^) (Figure 4D). Finally, the RCOR1 C-terminus (RCOR1^C^) is situated near the HDAC1-end of the CoREST structure, above RCOR1^SANT2^ (Figure 4D). HDAC1 is localized precisely, forming interactions with the KDM1A^TOWER^ domain, RCOR1^N^, and RCOR1^SANT2^, apart from RCOR1^ELM-SANT1^ to which it is bound (Figure 4D-E). The HDAC1 C-terminus (HDAC1^C^) appears to be localized poorly (Figure 4D-E). While our modeled CoREST architecture is overall consistent with a previous report ^47^, we also report several newly predicted protein-protein contact regions that include regions not covered by existing structural information (Figure 4F, Table S2).

### Modeling the intrinsically disordered C-Terminal domain of HDAC1

We next sought to better understand how HDAC1 and its intrinsically disordered CTD may be interacting with specific proteins within its protein interaction network (Table S1). To begin, we used AlphaFold3 ^40^ multimer to predict dimeric (Fig. S8) and trimeric (Fig. S9) assemblies of HDAC1 protein with RCOR1, SIN3A, MBD3, and MTA1 proteins without any other information than the protein sequences. Lysine-lysine Euclidian distances plots and residue contact maps were calculated for AlphaFold-only dimers of RCOR1:HDAC1, SIN3A:HDAC1, MBD3:HDAC1, and MTA1:HDAC1 (Fig. S8A-K). In each case, the AlphaFold models of each dimer show significant amounts of disorder remaining and the CTD of HDAC1 remains largely unstructured (Fig. S8C, F, I, and L). Since both MBD3 and MTA1 are both members of the NurRD complex and they both crosslinked to HDAC1, we created an AlphaFold-only model of the HDAC1:MDB3:MTA1 trimer (Fig. S9). Here, the disorder in the system decreased with improved modeling of the CTD of HDAC1 with contacts observed with both MDB3 and MTA1 (Fig. S9A-D), although large loops remained unstructured in all three proteins (Fig. S9 E-F).

We next used an integrative structural modeling (ISM) approach combining AlphaFold3, crosslinking mass spectrometry data, and molecular docking ^61^ to model dimeric (Figure 5) assemblies of HDAC1 protein in the presence of RCOR1, SIN3A, MBD3, and MTA1 proteins. In each of the 4 dimers, the HDAC1 CTD folded into a largely alpha helical structure (Fig. 5C, F, I, L). However, in the ISM models of the RCOR1:HDAC1, SIN3A:HDAC1, MBD3:HDAC1, and MTA1:HDAC1 dimers, HDAC1 CTD showed limited to no contact with the individual protein modeled with it, as shown in the lysine-lysine distance and residue contact maps (Figure 5A-B, D-E, G-H, J-K) We then built an ISM model of the HDAC1:MBD3:MTA1 trimer based on XL-MS data that resulted in an ordered and compact model with the HDAC1 CTD forming an alpha helical structure (Figure 6). A comparison of the dimers and trimers modeled using an AlphaFold-only approach and the integrative structural modeling approach (Fig. S10A-F) further demonstrated that the ISM approach was able to fully model a folded HDAC1 CTD within the dimers and trimer whereas the AlphaFold-only approach was not.

**Figure 5.**
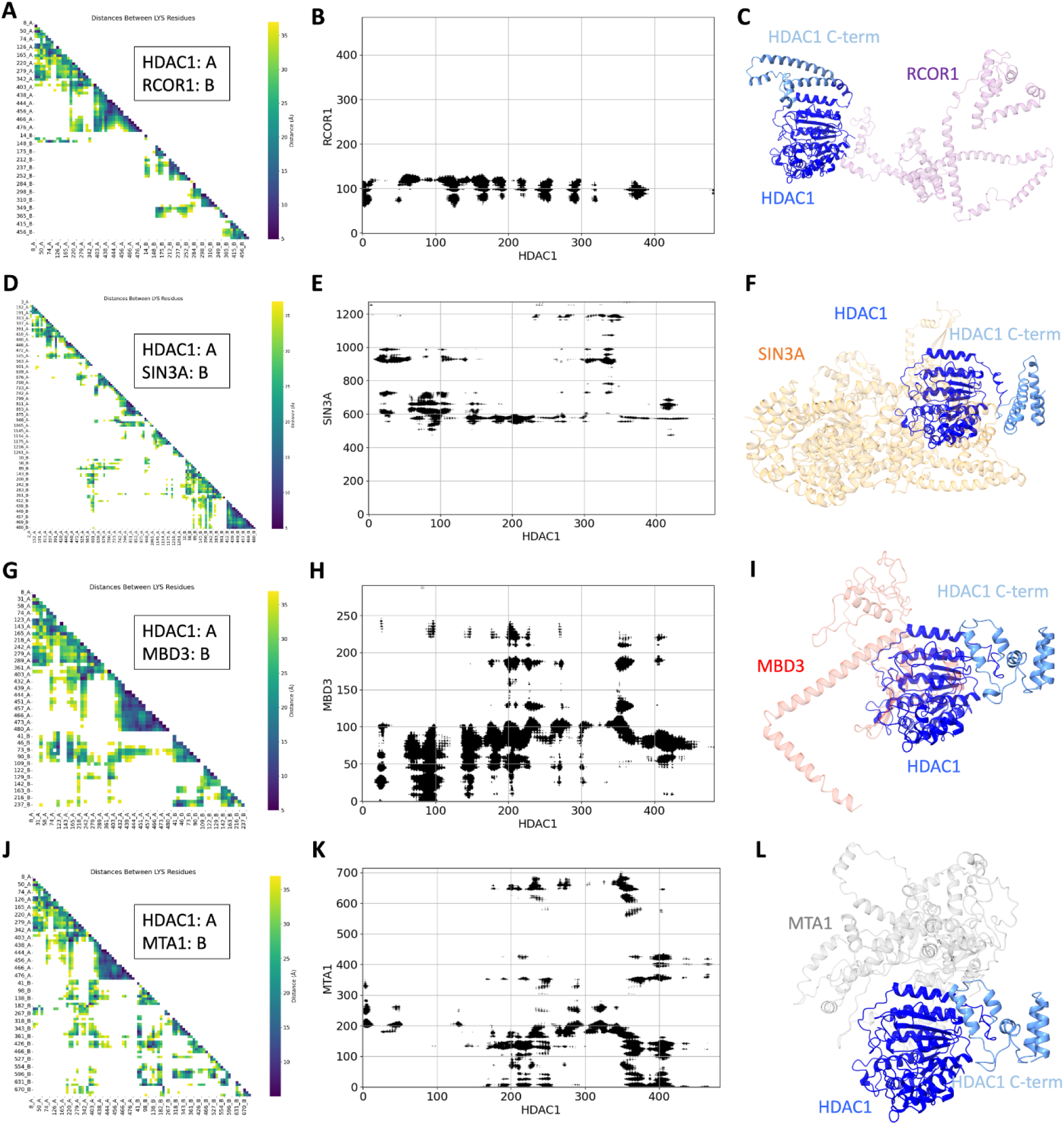
Interaction and structural analysis of the integrative dimeric complexes. (A-C) HDAC1/RCOR1 complex; (D-F) HDAC1/SIN3A complex; (G-I) HDAC1/MBD3 complex; (J-L) HDAC1/MTA1 complex. (A, D, G, and J) Lysine-lysine interaction maps for the different dimeric complexes, showing key lysine interactions within each complex. (B, E, H, and K) Residue contact maps illustrating the positions of interacting residues for each dimer. (C, F, I, and L) 3D visualization of the different dimers, providing a comparative view of their structural arrangement, therefore, highlighting the close proximity in conformations of the HDAC1 in these complexes. The multiple conformations of the HDAC1 C-termini are shown in light blue.

**Figure 6.**
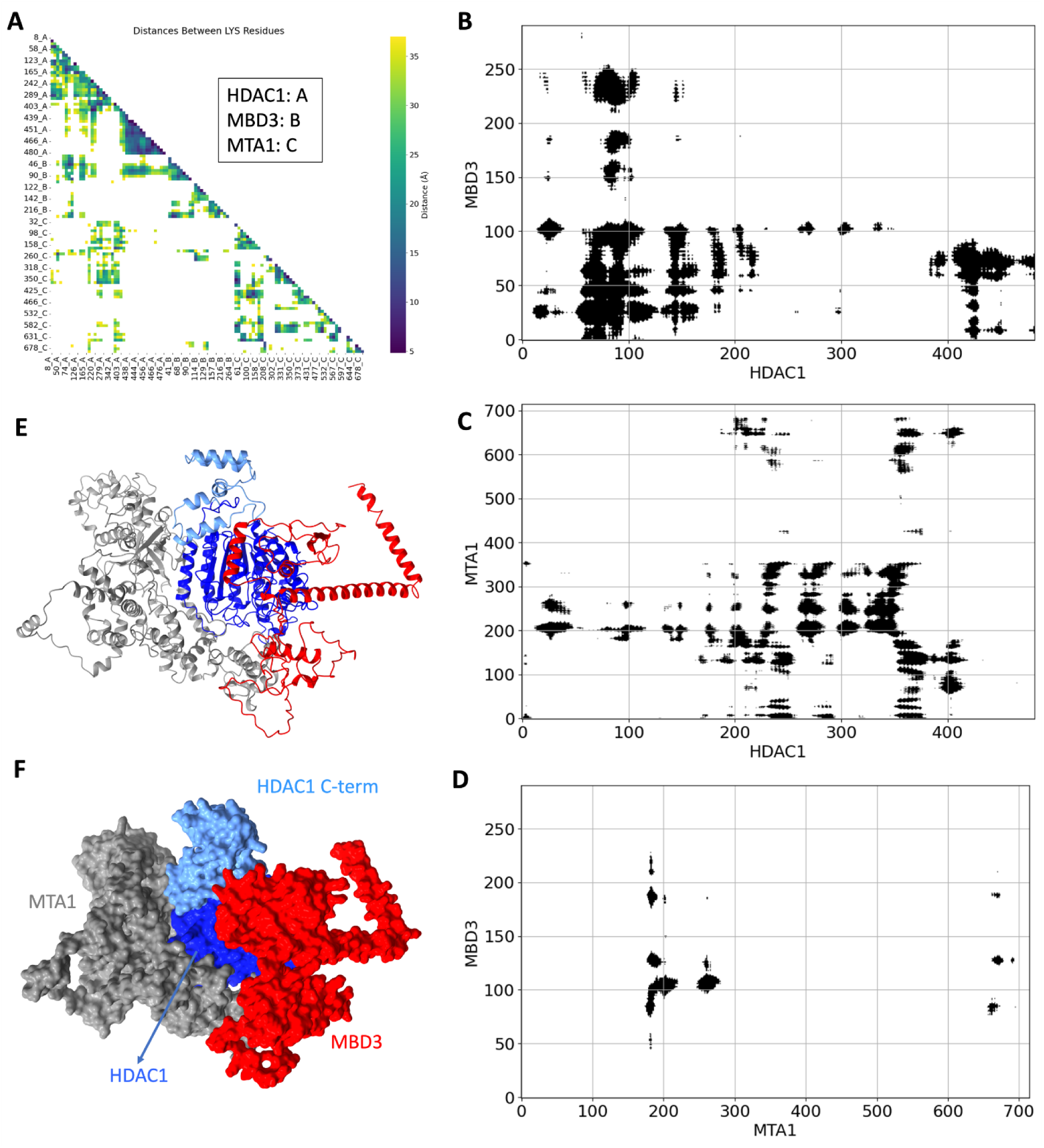
Structural and interaction analysis of the integrative trimeric HDAC1/MBD3/MTA1 Complex. (A) Lysine-lysine interaction maps for the HDAC1/MBD3/MTA1 trimer, illustrating key lysine residues between all three components. (B-D) Residue contact maps showing the interactions between HDAC1 and MBD3 (B), HDAC1 and MTA1 (C), or MTA1 and MBD3 (D) within the trimeric complex. (E, F) 3D visualization of the HDAC1/MBD3/MTA1 trimer and its surface representation (F) showcasing the spatial arrangement of each subunit within the complex to trigger a unique conformation of the HDAC1 (as well as the C-terminal domain of HDAC1 shown in light blue) in the context of the trimeric complex.

### Integrative structural modeling of Intrinsically Disordered Domains in a NuRD Subcomplex

Since the XL-MS guided approach was able to model the IDR within HDAC1 CTD, we next sought to model additional IDRs in the members of HDAC1-containing complexes. Crosslinks within the NuRD complex were the most abundant in the XL-MS dataset (Table S1), which we then used to analyze a larger subcomplex. From this dataset, we used 47 unique intramolecular and intermolecular crosslinks found within and between HDAC1, MBD3, MTA1, GATAD2B and RBBP4 (Table S3). In addition to the IDR within HDAC1 CTD, MBD3 has 1 IDR (from amino acids (AAs) 254 to 291), MTA1 has 2 IDRs (AAs 435-460 and 673-715), and GATAD2B has 2 IDRs (AAs 62-123 and 213-235), as called by UniProt ^10^ (Table S4). Within each of these IDRs, AlphaFold models potential pre-structured motifs (PreSMos), which are transient and locally ordered structural elements that are primed for binding ^62,63^ (Figure S11A).

Next, we used the ISM approach to model the HDAC1:MBD3:MTA1:GATAD2B:RBBP4 subcomplex. The disordered regions of each individual protein formed a series of structural elements largely dominated by alpha helices (Fig. S11B). The initial ISM model of this subcomplex (Figure S12A) successfully satisfied 87% of the experimentally observed crosslinks (Table S3), indicating strong consistency with crosslinking mass spectrometry data. However, the presence of 6 unmatched crosslinks (Table S3) suggested potential conformational variability or alternative structural arrangements. These unmatched crosslinks were subsequently used to generate an alternative model (Figure S12B). The comparison of these 2 models (Figure S12C) highlights regions of potential conformational flexibility within the NuRD complex.

Each protein within the subcomplex was distinctly color-coded to visualize their key domains and regions of functional importance (Fig. 7A), and a detailed description of the domains and secondary structures within each protein is reported in Table S4. To begin, in HDAC1, we focused on the C-terminal region and the metal-binding catalytic site, both of which are critical for its histone deacetylase activity. The C-terminal region of HDAC1, which initially displays IDR characteristics in its monomeric form (Fig. S11A and Table S4), folds into an ordered alpha helical structure suggesting potential flexibility in protein interactions and regulation. MBD3 is shown with its methyl-CpG-binding domain (MBD) and IDRs (Fig. 7A). MTA1 is highlighted with its ELM2 and SANT domains, along with surrounding IDRs and GATA-type zinc finger domains (Fig. 7A). The ELM2 and SANT domains are likely important for mediating protein-protein interactions, particularly in recruiting HDAC1 and facilitating the assembly of deacetylase modules (Fig. 7B-C). Similarly, GATAD2B is represented with its IDRs, its CR1 and CR2 regions, and GATA-type domain (Fig. 7A). The organization of each of the individual proteins is then shown in ribbon (Figure 7B) and in space filled (Figure 7C) forms. Individual proteins and domains within proteins are color coded to highlight the domains listed above in each protein in the context of the HDAC1:MBD3:MTA1:GATAD2B:RBBP4 subcomplex where a total of six IDRs are modeled. The presence of IDRs coupled with pre-structured motifs across these proteins, including the C-terminal tail of HDAC1, over 50% of the GATAD2B sequence, and IDRs interspersed with the ELM2 and SANT domains of MTA1 (Fig. S11A and Table S4) appear to provide the structural flexibility necessary for conformational selection coupled with an induced fit mechanism, where binding partners are accommodated through dynamic structural rearrangements ^17^.

**Figure 7.**
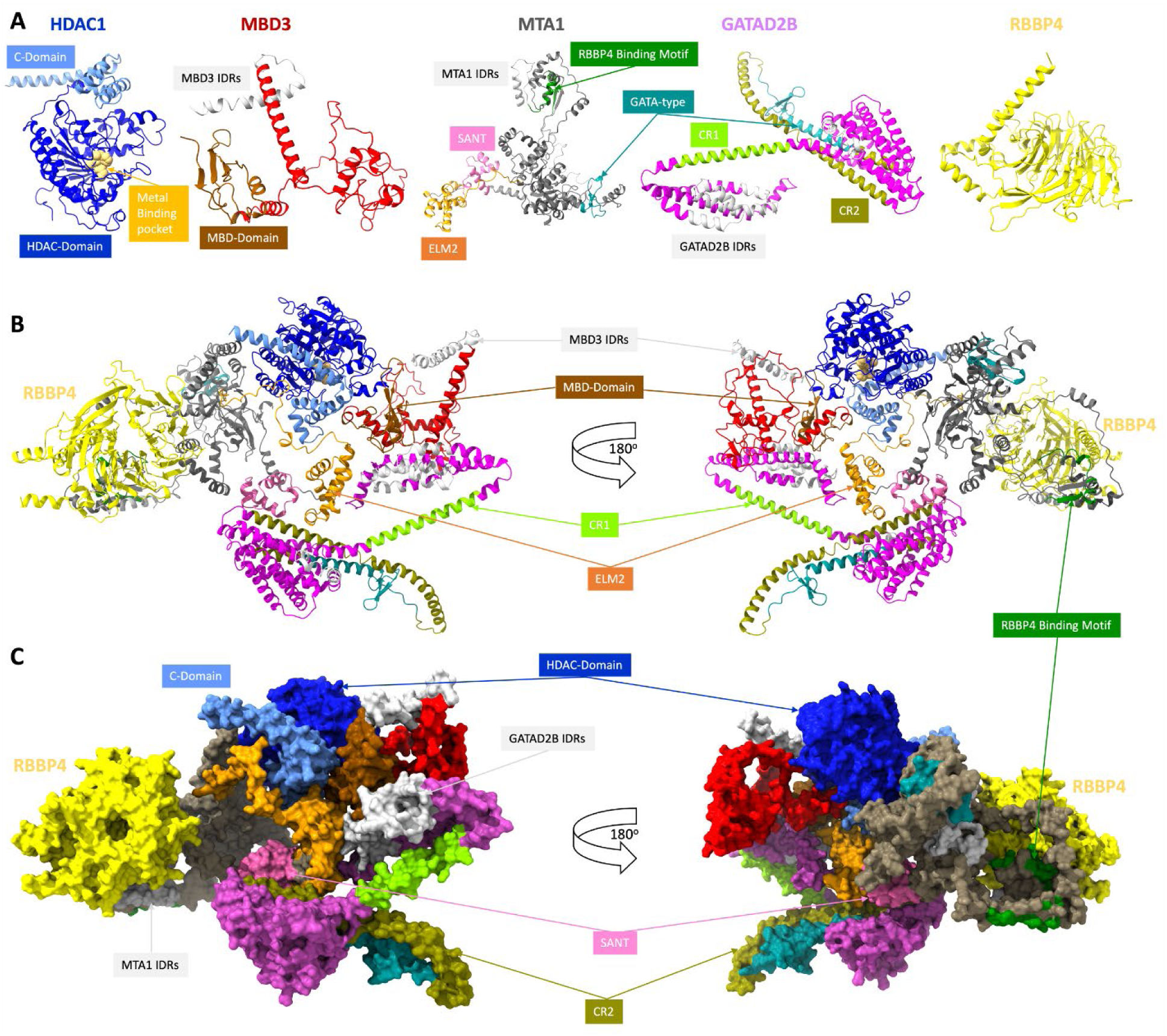
Detailed 3D structural representation of the NuRD sub-complex formed by HDAC1:MBD3:MTA1:GATAD2B:RBBP4. **(A)** Each protein component is distinctly color-coded to highlight specific functional domains and regions critical for complex assembly and function. For HDAC1, the C-terminal region and the metal-binding catalytic site are emphasized, revealing its enzymatic core. MBD3 is shown with its MBD domain and extensive intrinsically disordered regions (IDRs), reflecting its dual role in DNA interaction and dynamic scaffolding. MTA1 is depicted with its ELM2 and SANT domains, which are central to mediating protein-protein interactions, as well as its IDRs and GATA-type zinc finger domains, contributing to its regulatory flexibility. GATAD2B is illustrated with its IDRs, CR1 and CR2 (conserved regions), and a GATA-type domain, which facilitate both structural integration and functional specificity within the NuRD complex. **(B, C)** The ELM2 and SANT domains of MTA1 are particularly prominent, as they serve as critical hubs for recruiting and stabilizing interactions among the subunits, thereby orchestrating the assembly and functionality of the complex. RBBP4 is shown in association with the complex, completing the core structural framework. This comprehensive visualization underscores the modular architecture and cooperative nature of the complex, essential for its role in chromatin remodeling and transcriptional regulation.

## Discussion

In this study, we used HEK293 cells that stably expressed HDAC1-Halo and HDAC2-Halo to isolate HDAC1- and HDAC2-containing protein complexes that were analyzed by XL-MS. In doing so, we collected a large dataset of direct protein-protein interactions and built integrative structural models of subcomplexes of these endogenous chromatin remodeling complexes including NuRD ^4,35,36^, SIN3A^4,37^, and CoREST^4,38,39^. To begin, we deployed Bayesian integrative structural modeling via the Integrative Modeling Platform, which combines structural and biochemical data at several scales to build detailed models of complexes ^35,64^.

We first modeled the NuRD complex. Our results, which are validated by a previously published NuRD integrative model ^35^, allowed us to extend our XL-MS data for use in modeling the CoREST and SIN3 complexes. Understanding the structure of co-repressor complexes is critical for understanding their function. Based on the structures of the three HDAC co-repressor complexes obtained in the current study, we attempted to elucidate the design elements of a co-repressor complex. The four general components of a co-repressor complex include a scaffolding protein, which functions to assemble other subunits, one or more enzymes with chromatin-modifying activity (e.g., deacetylase, demethylase) and one or more DNA-binding domains, and transcription factor-binding domains.

First, the scaffolding protein, which consists of MTA1 in NuRD, SIN3A in SIN3A, and RCOR1 in CoREST, plays an important role in the assembly of the complex as well as mediates recruitment to the target site. The interaction of HDAC with the scaffolding protein could be tightly coupled, as in the case of complexes where an ELM-SANT domain of the scaffold interacts with HDAC, as in CoREST, MIER, and NuRD. For example, MTA1 in NuRD provides the scaffold for HDAC while also dimerizing to include two HDACs. Alternatively, the scaffold could be loosely coupled with HDAC and not involve an ELM-SANT domain, e.g., SIN3A. Here, the scaffolding and dimerization functions are split between SIN3A and SUDS3, respectively. This latter scaffold may require additional proteins, such as SAP30 for stabilizing the complex^37^. Second, each complex must bind to specific transcription factors to be recruited to the target site, mediated by specific subunits. RBBPs in NuRD, RCOR1 in CoREST, and SIN3A and SAP30 in SIN3A complex associate with transcription factors^23,47,65^. DNA-binding domain containing proteins may also help recruiting the complex to the target site. In NuRD, MBD3 contains binding sites for methylated and hemi-methylated DNA. Other potential DNA-binding domains also exist: for instance, MTA1^BAH^, MTA1^SANT2^, MTA1^ZFD^ in NuRD, RCOR1^SANT2^ in CoREST, and SAP30^ZFD^, SUDS3 in SIN3A complex.^66^. Third, corepressor complexes may harbor one or more chromatin-modifying proteins. SIN3A contains only a deacetylase with HDACs 1 and 2. However, both NuRD and CoREST contain other enzymes. NuRD consists of a deacetylase (HDAC1/2) and an ATP-dependent chromatin remodeler (CHD3/4). CoREST contains a deacetylase (HDAC1/2) and demethylase (KDM1A/LSD1)^47^. These corepressor complexes may either act on one or more nucleosomes, or even on two different targets within the same nucleosome simultaneously. CoREST harbors a single copy of the two chromatin modifier enzymes and is likely that these enzymes act in a concerted manner, with one enzyme acting on a nucleosome at a time^47^. NuRD and SIN3A on the other hand contain more than one copy of HDAC1/2, which may allow them to act on multiple sites. For SIN3A, SUDS3 may allow dimerization of the entire complex which may make it possible to act on adjacent nucleosomes simultaneously.

We next sought to investigate how HDAC1 might fold into each complex with a particular focus on the CTD IDR, which is currently absent from all HDAC1 structures ^10^ and is presented as a long and disordered string in AlphaFold ^40^. We used our XL-MS data, AlphaFold ^40^, and HADDOCK ^41^ to model the complete sequence of HDAC1 with complete sequences of RCOR1, SIN3A, MBD3, and MTA1, which were all proteins crosslinked to HDAC1 in our dataset. To begin, the comparison of HDAC1 in complex with RCOR1 (CoREST complex) and SIN3A (SIN3A complex) provides an example of how different protein complexes can alter the folding of HDAC1 in distinct ways. Although the central, conserved region of HDAC1 remains largely unaffected by the presence of either RCOR1 or SIN3A, our results show that the IDR C-terminal region exhibits marked differences in flexibility (Fig. 5A-F). In the presence of RCOR1, HDAC1 adopts a more flexible conformation, particularly in its C-terminal tail, which is consistent with previous observations suggesting that the CoREST complex is associated with a more dynamic, regulatory role in chromatin remodeling ^47^. This increased flexibility could be essential for the functional interactions required for gene repression by HDAC1/RCOR1. Conversely, when HDAC1 is bound to SIN3A, the C-terminal region adopts a more rigid conformation (Fig. 5F). SIN3A, a component of the SIN3A complex, appears to stabilize the C-terminal tail of HDAC1 (Fig. 5F), potentially favoring a more fixed, structured state ^65,67,68^. This structural difference may reflect the distinct functional roles of the two complexes—CoREST and SIN3A—both of which regulate transcriptional repression, but likely in different cellular contexts and with divergent mechanisms ^1^. The fact that the middle region of HDAC1 remains largely unchanged in both complexes underscore the conservation of its core enzymatic function, despite variations in complex composition.

In contrast to the differential effects observed with RCOR1 and SIN3A, the binding of HDAC1 to either MBD3 or MTA1—two components of the NuRD complex ^26,69^—produced similar structural outcomes (Fig. 5 G-L). Both HDAC1/MBD3 and HDAC1/MTA1 complexes showed minimal variation in their overall structure, with the middle and C-terminal regions of HDAC1 adopting near-identical conformations (Fig. 5 I, L). This similarity is likely because MBD3 and MTA1 are part of the same complex, and their interactions with HDAC1 are more synergistic and less context-dependent than those of RCOR1 and SIN3A. When the HDAC1/MBD3/MTA1 complex was modeled as a trimer, rather than as separate dimers, the overall structure became more compact (Fig. 6). This reinforces the importance of considering multi-protein complexes in structural modeling, as the dimeric form of HDAC1/MBD3 or HDAC1/MTA1 (Fig. 5 G-L) alone does not fully capture the conformational state of HDAC1 in a cellular environment where these proteins function together in larger complexes (Fig. 6).

In all cases where dimers or trimers were modeled the CTD IDR of HDAC1 folded into a compact alpha helical structure. With this ability to model IDRs, we then sought to build a model of a NuRD subcomplex to investigate the behavior of multiple IDRs in a system. The 3D model of the NuRD subcomplex, as illustrated in Sup. Video 1, suggests a symmetric and modular assembly within the NuRD complex, involving both monomeric and dimeric interfaces, which may be essential for the dynamic function of the full NuRD complex ^12,27,70^. The color-coding of key domains of the model formed by HDAC1:MBD3:MTA1:GATAD2B:RBBP4, such as HDAC1 C-terminal and metal-binding regions, MBD3 MBD domain and IDRs, MTA1 ELM2, SANT, and GATA-type domains, and GATAD2B IDRs as well as its CR1 and CR2 conserved regions highlights their structural and functional roles (Fig. 7) ^8,21,23,70–72^. The ELM2 and SANT domains of MTA1 play a central role in complex assembly by stabilizing interactions with HDAC1 and contributing not only to the recruitment of other subunits of the sub-complex, but also to the overall stability of the structure ^71^.

An important insight from this model is the importance of intrinsically disordered regions, which are prominent in HDAC1, MTA1, MBD3, and GATAD2B and likely contribute to structural flexibility and adaptive binding via a conformational selection/induced fit mechanism ^21,62,73^. The IDRs in HDAC1, MTA1, MBD3, and GATAD2B contain pre-structured motifs, which serve as conserved molecular recognition elements critical for specificity and regulation ^8,23,73^. Our model supports the view of NuRD as a dynamic, compositionally flexible complex capable of forming distinct assemblies, such as a deacetylation-only module or a full chromatin remodeling complex, depending on cellular context ^24^.

Our results highlight the importance of the cellular environment in determining the conformation of HDAC1 ^1–3,74^. The structural plasticity of HDAC1, influenced by the presence of distinct protein partners such as RCOR1, SIN3A, MBD3, and MTA1, suggests that HDAC1 function is tightly regulated by its interaction network. While the core, conserved region of HDAC1 may be primarily responsible for its catalytic activity, the variable flexibility of the N-and C-terminal regions appear to play a key role in modulating its interactions with other proteins, potentially affecting the stability and specificity of HDAC1-containing complexes. Additionally, this study also underscores the need for more complex structural models of HDAC1 that incorporate multiple interacting proteins, rather than relying on isolated dimers or simplified configurations. The structural differences we observed between HDAC1-containing dimers or trimers further suggest that protein-protein interactions are crucial for stabilizing the protein in a conformation that is relevant to its biological function. Given that HDAC1 is involved in diverse processes, including transcriptional regulation, chromatin remodeling, and cellular signaling ^1^, understanding how its structure is modulated by different complexes will provide important insights into its multifaceted role in the cell.

## Acknowledgements

The authors would like to thank Vinothkumar K.R., Shreyas Arvindekar, Muskaan Jindal, Omkar Golatkar, and Mubashira K.P. for helpful comments on the manuscript. The use of NCBS institutional computational and storage facilities is gratefully acknowledged. S.V. acknowledges support from the Department of Atomic Energy, Government of India, through grant DAE TIFR RTI 4006 and from the Department of Biotechnology, Government of India grant BT/PR40323/BTIS/137/78/2023. In addition, research reported in this publication was supported by the Stowers Institute for Medical Research and the National Institute of General Medical Sciences of the National Institutes of Health under award numbers F31GM131536 (C.G.K.), T32GM138077 (J.C.), R35GM118068 (J.L.W.), and R35GM145240 (M.P.W.). The content is solely the responsibility of the authors and does not necessarily represent the official views of the National Institutes of Health.

## Author Contributions

Conceptualization: JN, KM, SV, MPW

Investigation: JN, KM, RZ, CGK, JC, YZ

Formal Analysis: JN, KM, CGK, YZ

Resources: JN, KM, CGK, YZ, JT, LF, YZ, SV, MPW

Software: JN, KM, SV, MPW Data Curation: JN, KM, LF

Writing: JN, KM, RZ, LF, SV, MPW

Supervision: LF, JLW, SV, MPW

Funding acquisition: CGK, JLW, SV, MPW

## Data and Materials availability

Mass spectrometry data may be accessed through Proteome Xchange (PXD074062) via the MassIVE ftp repository (massive.ucsd.edu) using MSV000100718 as username and “HDAC-ISM_2026” as password. After publication, original mass spectrometry data underlying this manuscript may also be accessed from the Stowers Original Data Repository at https://www.stowers.org/research/publications/LIBPB-2492. Files containing the input data, scripts, and output results are publicly available at https://github.com/isblab/hdac. The input data, scripts, output results, and the final integrative structures have been deposited at Zenodo (DOI: 10.5281/zenodo.11056108). The integrative models for the NuRD, CoREST, and SIN3A complexes will be deposited to wwPDB.

## STAR METHODS

### Materials

Magne^®^ HaloTag^®^ magnetic affinity beads (G7281), Rabbit anti-HaloTag^®^ polyclonal antibody (G9281), Sequencing Grade Modified Trypsin (V5111), Mammalian Lysis Buffer (G938A), and Protease Inhibitor Cocktail (G6521) were purchased from Promega. HEK293T cells (ATCC1CRL-11268™) were purchased from American Type Culture Collection (ATCC). Flp-In™-293 cells (AHO1112) and AcTEV protease (#12575015) were purchased from Invitrogen. Opti-MEM™ Reduced Serum Medium (31985062), Pierce™ BCA Protein Assay Kit (23225), and GlutaMAX™ Supplement (35050061) were purchased from ThermoFisher Scientific. Salt Active Nuclease (HL-SAN, 70910-202) was purchased from ArcticZymes. Disuccinimidyl sulfoxide (DSSO, 9002863) was purchased from Cayman Chemical. Fetal Bovine Serum (FBS, PS-FB1) was purchased from PEAK^®^ Serum. Dulbecco’s Modified Eagle Medium (DMEM, 10-013-CV) was purchased from Corning. Hygromycin B (H-1012-PBS) was purchased from AG Scientific.

### Affinity Purification Mass Spectrometry of HDAC1-Halo and HDAC2-Halo

Whole cell extracts containing the Halo-tagged bait protein prepared from Flp-In™-293 cell lines were prepared as described previously ^37^. Briefly, for each replicate, 3 confluent 850 cm^2^ culture vessels of Flp-In™-293 cells stably expressing HDAC1-Halo and HDAC2-Halo (for XL-MS) were harvested, washed twice with ice-cold PBS, incubated at −80°C until frozen, thawed, and resuspended in ice-cold lysis buffer containing: 20 mM HEPES (pH 7.5), 1.5 mM MgCl2, 0.42 M NaCl, 10 mM KCl, 0.2% Triton X-100, 0.5 mM DTT, 0.1 mM benzamidine HCL, 55 µM phenanthroline, 10 µM bestatin, 20 µM leupeptin, 5 µM pepstatin A, 1mM PMSF, and 500 units SAN. Following Dounce homogenization, extracts were incubated for 2 hours at 4°C and centrifuged 40,000 × g for 30 minutes at 4°C. The salt concentration was adjusted to 0.3 M NaCl by adding ice-cold buffer containing: 10 mM HEPES (pH 7.5), 1.5 mM MgCl2, 10 mM KCl, 0.5 mM DTT, 0.1 mM benzamidine HCl, 55 µM phenanthroline, 10 µM bestatin, 20 µM leupeptin, 5 µM pepstatin A, and 1mM PMSF. Lysates were centrifuged at 40,000 × g for 30 minutes at 4°C for further analysis.

Protein extracts were next incubated with Magne^®^ HaloTag^®^ beads prepared from 200 µl Magne™ HaloTag^®^ bead slurry and crosslinked with disuccinimidyl sulfoxide (DSSO, Cayman Chemical Company) MS-cleavable crosslinker as previously described ^32,33,37^. Briefly, DSSO was added to samples while the proteins were immobilized on beads to a final concentration of 5 mM. Samples were incubated at room temperature for 40 min. Reactions were quenched with the addition of NH4CO3 to a final concentration of 50 mM and samples were incubated an additional 15 min. Bound proteins were eluted by incubating the beads with AcTEV™ Protease (Invitrogen) overnight at 25°C.

### Crosslinking Mass Spectrometry and Data Analysis

Digested peptides were analyzed with an Orbitrap Fusion™ Lumos™ (Thermo Fisher) coupled to a Dionex U3000 HPLC using previously described data acquisition settings^31,37^. From each .RAW file, the XlinkX node in Proteome Discoverer v2.4 (Thermo Fisher) ^75^ was used to identify MS2 fragmentation scans with reporter ions characteristic of DSSO crosslinked peptides and Sequest HT was used to identify linear peptides. Peak lists were searched against a human proteome database (Genome Reference Consortium Human Build 38 patch release 13) containing 44519 unique protein sequences, 426 of which were contaminant proteins. The database was searched for fully tryptic peptides, allowing for a maximum of 2 missed cleavages and a minimum peptide length of 5 amino acids. Precursor mass tolerance, FTMS fragment mass tolerance, and ITMS Fragment tolerance, were set to 10 ppm, 20 ppm, and 0.5 Da, respectively. Xlink Validator FDR threshold was set to 0.01. Searches were performed with a static modification of +57.021 Da on cysteine, a dynamic modification of +15.995 Da on methionine residues, a dynamic modification of +176.014 Da on lysine residues (water-quenched hydrolyzed DSSO monoadduct), and a dynamic modification of 279.078 Da on lysine residues (Tris-quenched DSSO monoadduct).

### Integrative Modeling Platform

Integrative structure determination of three HDAC1/2-containing co-repressor complexes (NuRD, CoREST, and Sin3A) proceeded through four stages ^35,64^. The modeling protocol (i.e., stages 2, 3, and 4) was scripted using the Python Modeling Interface (PMI) package, a library for modeling macromolecular complexes based on open-source Integrative Modeling Platform (IMP) package, version 2.16.0 and v.23.0 (https://integrativemodeling.org) ^64^. The current procedure is an updated version of previously described protocols ^34,35^. Files containing the input data, scripts, and output results are publicly available at https://github.com/isblab/hdac. All integrative models will be deposited in the wwPDB.

#### Stage 1: Gathering Data

*NuRD–*For each subunit, the paralog with the most crosslinks was chosen for modeling. Our NuRD model included full-length MTA1, HDAC1, RBBP4, MBD3, and GATAD2B^1-281^, based on the presence of crosslinks for each protein/domain (Figure 2A). Known atomic structures were used for the MTA1-HDAC1 dimer, MTA1^R1^ and MTA1^R2^ domains in complex with RBBP4, and MBD domain of MBD3 (Fig. S2A-B) ^8,50,51^. The MTA1^BAH^ domain, MTA1^H^, MTA1^ZF^, and MBD3^CC^-GATAD2B^CC^ structures were homology-modeled based on the structures of related templates (Fig. S2A-B) ^35,36,53,76^. AlphaFold2 was used to obtain structures for MTA1, MBD3, GATAD2B for which no structure or homology model was available (Fig. S2A-B) ^40^. Regions of high confidence (>70 pLDDT and <5 PAE) and at least 20 residues in length were used. The shape of the complex was based on cryo-EM map EMD-22895 (14 Å) ^36^ (Fig. S2B). Chemical crosslinks informed the relative localization of the NuRD subunits. For modeling the NuRD complex, 73 DSSO (Bis(2,5-dioxopyrrolidin-1-yl) 3,3′-sulfinyldipropionate, Bis-(propionic acid NHS ester)-sulfoxide) crosslinks were used (Fig. S2C). Crosslinks from paralogs MTA2, MTA3 were mapped to MTA1; RBBP7 to RBBP4; MBD2 to MBD3; GATAD2A to GATAD2B using sequence alignments. The subunit stoichiometry (Fig. S2D) was based on DIA-MS and SEC-MALS experiments ^36^.

*SIN3A*—For each subunit, the paralog with the most crosslinks was chosen for modeling (Figure 3). For SIN3A, our model includes SIN3A^456-831^ (SIN3A^PAH3^, SIN3A^HID^) and full-length HDAC1, SAP30, and SUDS3. Known atomic structures were used for SAP30^ZFD^ and HDAC1 (Fig. S4A-B) ^12,77^. SAP30^SID^-SIN3A^PAH3^, SUDS3^SID^ -SIN3A^HID^, and SUDS3^CC^ - were homology modelled based on structures of related templates (Fig. S4A-B) ^78,79^. AlphaFold2 was used to obtain structures for regions of SIN3A, SUDS3 for which no structure or homology model was available, with structures pre-processed as described above (Fig. S4A-B) ^40^. Interacting residues/domains for SAP30-HDAC1^66^ and HDAC1-SIN3A ^78^ were obtained via co-immunoprecipitation and SEC-MALS, respectively. These were modelled as minimum pair distance restraints which anchor the interacting residues/domains together. Chemical crosslinks informed the relative localization of the Sin3A subunits. For modelling the SIN3A complex, 32 DSSO crosslinks were used (Fig. S4C). Crosslinks from SAP30L were mapped to SAP30 using sequence alignment. Stoichiometry of SIN3A was assumed based on previous studies ^31,78^ (Fig. S4D).

*CoREST*—For each subunit, the paralog with the most crosslinks was chosen for modeling (Figure 4). For CoREST our model includes full-length RCOR1, HDAC1, KDM1A (Fig. S6A-B). A known atomic structure was used for KDM1A-RCOR1 (Fig. S6A-B) ^80,81^. RCOR1^ELM2-SANT1^-HDAC1 was homology modelled using the MTA1-HDAC1 structure as template (Fig. S6A-B) ^8^. The shape of the complex was based on glutaraldehyde crosslinked cryo-EM map EMD-10627 (17.5 Å) (Fig. S6B) ^47^. Chemical crosslinks informed the relative localization of the CoREST subunits. For modelling the CoREST complex, 11 DSSO crosslinks were used (Fig. S6C). Crosslinks from RCOR2 were mapped to RCOR1 using sequence alignments. The stoichiometry of subunits was based on SEC experiments (Fig. S6D) ^39^.

#### Stage 2: Representing the system and translating data into spatial restraints

The domains with known atomic structures were represented in a multi-scale manner with 1 and 10 residues per bead to maximize computational efficiency ^82^. These domains were modelled as rigid bodies where the relative distances between beads is constrained during sampling. In contrast, domains without known structure were coarse-grained at 30 residues per bead (10 residues per bead for flexible regions of SUDS3 173-204) and modeled as flexible strings of beads. The representations of subunits for each complex are shown in Figures S2A, S4A, and S6A.

We next encoded the spatial restraints into a scoring function based on the information gathered in Stage 1, as follows:

1. Crosslink restraints: The Bayesian crosslinks restraint was used to restrain the distances spanned by the crosslinked residues. The restraint accounts for ambiguity (multiple copies of a subunit) via a compound likelihood term that considers multiple residue pairs assigned to an individual crosslink.
2. EM restraints: The Bayesian EM density restraint was used to restrain the shape of the modelled complexes and was based on the cross-correlation between the Gaussian Mixture Model (GMM) representations of the subunits and the GMM representation of the corresponding cryo-EM density maps ^83^.
3. Minimum Pair Distance Binding restraint: Distance restraints were used to restrain the interacting regions obtained from coimmunoprecipitation. The restraint was encoded as a harmonic upper bound on the minimum distance between bead pairs of interacting domains. Ambiguity (multiple copies) was considered by creating a separate restraint for each copy of a molecule.
4. Excluded volume restraints: The excluded volume restraints were applied to each bead, using the statistical relationship between the volume and the number of residues that it covered.
5. Sequence connectivity restraints: We applied the sequence connectivity restraints, using a harmonic upper distance bound on the distance between consecutive beads in a subunit, with a threshold distance equal to twice the sum of the radii of the two connected beads. The bead radius was calculated from the excluded volume of the corresponding bead, assuming standard protein density.

For Sin3A, we applied the crosslink restraint, distance binding restraint, connectivity, and excluded volume restraints. For the other two complexes, we used the EM restraint, crosslink restraint, connectivity, and excluded volume restraints.

#### Stage 3: Structural sampling to produce an ensemble of structures that satisfies the restraints

We aimed to maximize the precision at which the sampling of good-scoring solutions was exhaustive (Stage 4). The sampling runs relied on Gibbs sampling, based on the Replica Exchange Monte Carlo algorithm ^34^. The positions of the rigid bodies (domains with known structure) and flexible beads (domains with unknown structure) were sampled.

The initial positions of the flexible beads and rigid bodies in all complexes were randomized, with one exception. For CoREST, we were able to unambiguously dock the structure of the KDM1A-RCOR1 dimer in the EM map, with the help of the previous EM map (EMD-10627) ^47^. Hence, the position of the corresponding rigid body was fixed throughout. The Monte Carlo moves included random translations of individual beads in the flexible segments and random rotations and translations of rigid bodies and super rigid bodies. A model was saved every 10 Gibbs sampling steps, each consisting of a cycle of Monte Carlo steps that moved every bead and rigid body once. The sampling produced a total of 72 million NuRD integrative models, 36 million CoREST models, 120 million models for SIN3A monomer and dimer.

#### Stage 4: Analyzing and validating the ensemble of structures and data

The sampled models were analyzed to assess sampling exhaustiveness and estimate the precision of the structure, its consistency with input data and consistency with data not used in modelling. The structure was further validated by experiments based on the predictions from the models. We used the analysis and validation protocol published earlier ^34,35^. Assessment began with a test of the thoroughness of structural sampling, including structural clustering of the models, estimating model precision (A-D panels in Fig. S3, S5, S7), and visualizing the variability in the ensemble of structures using localization probability density maps (E-F panels in Fig. S3, S5, S7). The positional variation of a domain in an ensemble of superposed models can be visualized by the localization probability density map for the domain, which specifies the probability of a voxel (3D volume unit) being occupied by a bead in a set of superposed models. Regions of high and low precision were computed using PrISM and visualized on the cluster center bead model ^84^ (G panel in Fig. S3, S5, S7). All models and densities were visualized with UCSF Chimera and ChimeraX ^85^.

1. Determining good-scoring models: Starting from the millions of sampled models, first, we selected models obtained after score equilibration and clustered them based on the restraint scores ^34^. For further analysis, we considered 28,914 NuRD, 16,055 CoREST, and 29,602 SIN3A good-scoring models that satisfy the data restraints sufficiently well (A-B panels in Fig. S3, S5, S7).
2. Clustering and structure precision: We next assessed the sampling exhaustiveness and performed structural clustering ^34^. Integrative structure determination resulted in effectively a single cluster for all complexes, at a precision of 55 Å (NuRD), 12 Å (CoREST), and 33Å (SIN3A). The model precision is the bead RMSD from the cluster centroid model averaged over all models in the cluster (C-D panels in Fig. S3, S5, S7).
3. Fit to input information: The fit to crosslinks was computed by obtaining the percentage of satisfied crosslinks; a crosslink is satisfied by a cluster of models if the corresponding Cα-Cα distance in any model in the cluster is less than 35Å. The NuRD, CoREST, and SIN3A models satisfied over 90% of the respective crosslinks used (panel A-right in Figures 2-4). The agreement between the models and the corresponding EM maps was computed by calculating the cross-correlation of the combined localization probability densities of all subunits for the major cluster with the experimental EM map using the fitmap tool in UCSF Chimera ^85^. The cross-correlation for both NuRD and CoREST was higher than 0.85. The remainder of the restraints are harmonic, with a specified standard deviation. The cluster generally satisfied the excluded volume and sequence connectivity restraints. A restraint is satisfied by a cluster of models if the restrained distance in any model in the cluster (considering restraint ambiguity) is violated by less than 3 standard deviations, specified for the restraint. Most of the violations are small, and can be rationalized by local structural fluctuations, coarse-grained representation of the model, and/or finite structural sampling.
4. Protein-protein contact maps: Contact maps were created by computing the average distance between the beads across all the models in the major cluster for all protein pairs. We further selected contacts which are at an average distance less than 10 Å (Table S2).

### Initial Prediction of HDAC1 Structure Using AlphaFold

To predict the three-dimensional (3D) structure of HDAC1 in various protein complexes, we first used the AlphaFold3 protein structure prediction server ^40^. For the dimeric models, we input the sequence of HDAC1 along with the sequence of its interacting partner (RCOR1 or SIN3A) into AlphaFold3 to predict the corresponding structures. The AlphaFold3 server was run separately for HDAC1/RCOR1 and HDAC1/SIN3A, where each protein was modeled independently. For the HDAC1/MBD3 and HDAC1/MTA1 complexes, we input HDAC1 sequence along with the sequences of MBD3 and MTA1, respectively, to model the individual dimers. To generate the HDAC1/MBD3/MTA1 trimer, the sequences of HDAC1, MBD3, and MTA1 were combined, and the AlphaFold server was used to predict the structure of the entire trimeric complex. For each complex, multiple models were generated. We used a custom scoring approach to evaluate the compatibility of predicted models with XL-MS-derived crosslinking distance restraints (Cα–Cα distance ≤35 Å). Only complex structures that showed the highest number of satisfied crosslinks were retained for further analysis.

### Integrative Protein Complex Prediction via Guided Molecular Docking with HADDOCK

From the top-scoring complexes, we extracted individual proteins and assembled them into complexes using the HADDOCK2.4 ^86^ docking platform. Input structures were prepared using PDBTools ^41^ and custom-made python scripts. Distance restraints derived from XL-MS data were converted into ambiguous interaction restraints and used to guide the docking process. Multiple docking runs were performed, and the top-ranked clusters were selected based on HADDOCK scoring functions and the number of satisfied crosslinks. To improve the accuracy of the final model, an iterative refinement protocol was employed. In each round, the best-scoring complex was re-docked using updated restraints based on prior crosslink evaluation. This process was repeated until no further improvement in crosslink satisfaction was observed. Final complex models were visualized and analyzed using UCSF ChimeraX ^85^.

The predicted 3D structures were processed and analyzed using custom Python scripts. Structural comparisons were conducted by superimposing the predicted HDAC1 models from different complexes, focusing on the C-terminal and middle regions of the protein. This allowed for the identification of structural shifts or flexibility differences in HDAC1 when bound to RCOR1, SIN3A, MBD3, and MTA1. The key steps involved in the structure analysis involved lysine-lysine interactions that were predicted by identifying pairs of lysine residues in proximity within 30 Å of each other. In these interactions, we calculated the distance between lysine residues and generated interaction maps for each complex (RCOR1, SIN3A, MBD3, MTA1, and their corresponding HDAC1 interactions). Residue contact maps were also generated by calculating the Euclidean distances for each pair of interacting residues within 30 Å of each other. Contact maps were plotted to compare the density of residue-residue interactions in different complexes (HDAC1/RCOR1, HDAC1/SIN3A, HDAC1/MBD3, HDAC1/MTA1, and the HDAC1/MBD3/MTA1 trimer). The maps were then used to assess the extent of structural differences between the complexes. Finally, the 3D structures were visualized using the VMD package ^87^ to facilitate direct comparisons of HDAC1 in different complexes. The structures were superimposed based on the conserved HDAC1 domain to isolate differences in the multiple HDAC1s within different complexes. The superimpositions allowed us to evaluate structural variations in the C-terminal region in response to the presence of RCOR1 or SIN3A (CoREST and SIN3A complexes, respectively), as well as MBD3 and MTA1 (NuRD complex).

## References

1. Milazzo, G., Mercatelli, D., Di Muzio, G., Triboli, L., De Rosa, P., Perini, G., and Giorgi, F.M. (2020). Histone Deacetylases (HDACs): Evolution, Specificity, Role in Transcriptional Complexes, and Pharmacological Actionability. Genes (Basel) 11. 10.3390/genes11050556.

2. Zhou, C., Zhao, D., Wu, C., Wu, Z., Zhang, W., Chen, S., Zhao, X., and Wu, S. (2024). Role of histone deacetylase inhibitors in non-neoplastic diseases. Heliyon 10, e33997. 10.1016/j.heliyon.2024.e33997.

3. Patel, R., Modi, A., and Vekariya, H. (2024). Discovery and Development of HDAC Inhibitors: Approaches for the Treatment of Cancer a Mini-review. Curr Drug Discov Technol 21, e230224227378. 10.2174/0115701638286941240217102948.

4. Banks, C.A.S., Miah, S., Adams, M.K., Eubanks, C.G., Thornton, J.L., Florens, L., and Washburn, M.P. (2018). Differential HDAC1/2 network analysis reveals a role for prefoldin/CCT in HDAC1/2 complex assembly. Sci Rep 8, 13712. 10.1038/s41598-018-32009-w.

5. Gonneaud, A., Turgeon, N., Jones, C., Couture, C., Levesque, D., Boisvert, F.M., Boudreau, F., and Asselin, C. (2019). HDAC1 and HDAC2 independently regulate common and specific intrinsic responses in murine enteroids. Sci Rep 9, 5363. 10.1038/s41598-019-41842-6.

6. Winter, M., Moser, M.A., Meunier, D., Fischer, C., Machat, G., Mattes, K., Lichtenberger, B.M., Brunmeir, R., Weissmann, S., Murko, C., et al. (2013). Divergent roles of HDAC1 and HDAC2 in the regulation of epidermal development and tumorigenesis. EMBO J 32, 3176–3191. 10.1038/emboj.2013.243.

7. Dovey, O.M., Foster, C.T., and Cowley, S.M. (2010). Histone deacetylase 1 (HDAC1), but not HDAC2, controls embryonic stem cell differentiation. Proc Natl Acad Sci U S A 107, 8242–8247. 10.1073/pnas.1000478107.

8. Millard, C.J., Watson, P.J., Celardo, I., Gordiyenko, Y., Cowley, S.M., Robinson, C.V., Fairall, L., and Schwabe, J.W. (2013). Class I HDACs share a common mechanism of regulation by inositol phosphates. Mol Cell 51, 57–67. 10.1016/j.molcel.2013.05.020.

9. Bressi, J.C., Jennings, A.J., Skene, R., Wu, Y., Melkus, R., De Jong, R., O’Connell, S., Grimshaw, C.E., Navre, M., and Gangloff, A.R. (2010). Exploration of the HDAC2 foot pocket: Synthesis and SAR of substituted N-(2-aminophenyl)benzamides. Bioorg Med Chem Lett 20, 3142–3145. 10.1016/j.bmcl.2010.03.091.

10. UniProt, C. (2025). UniProt: the Universal Protein Knowledgebase in 2025. Nucleic Acids Res 53, D609–D617. 10.1093/nar/gkae1010.

11. Turnbull, R.E., Fairall, L., Saleh, A., Kelsall, E., Morris, K.L., Ragan, T.J., Savva, C.G., Chandru, A., Millard, C.J., Makarova, O.V., et al. (2020). The MiDAC histone deacetylase complex is essential for embryonic development and has a unique multivalent structure. Nat Commun 11, 3252. 10.1038/s41467-020-17078-8.

12. Millard, C.J., Fairall, L., Ragan, T.J., Savva, C.G., and Schwabe, J.W.R. (2020). The topology of chromatin-binding domains in the NuRD deacetylase complex. Nucleic Acids Res 48, 12972–12982. 10.1093/nar/gkaa1121.

13. Yeo, M.J.R., Zhang, O., Xie, X., Nam, E., Payne, N.C., Gosavi, P.M., Kwok, H.S., Iram, I., Lee, C., Li, J., et al. (2025). UM171 glues asymmetric CRL3-HDAC1/2 assembly to degrade CoREST corepressors. Nature 639, 232–240. 10.1038/s41586-024-08532-4.

14. Wan, M.S.M., Muhammad, R., Koliopoulos, M.G., Roumeliotis, T.I., Choudhary, J.S., and Alfieri, C. (2023). Mechanism of assembly, activation and lysine selection by the SIN3B histone deacetylase complex. Nat Commun 14, 2556. 10.1038/s41467-023-38276-0.

15. Chen, Z., Chi, G., Balo, T., Chen, X., Montes, B.R., Clifford, S.C., D’Angiolella, V., Szabo, T., Kiss, A., Novak, T., et al. (2025). Structural mimicry of UM171 and neomorphic cancer mutants co-opts E3 ligase KBTBD4 for HDAC1/2 recruitment. Nat Commun 16, 3144. 10.1038/s41467-025-58350-z.

16. Holehouse, A.S., and Kragelund, B.B. (2024). The molecular basis for cellular function of intrinsically disordered protein regions. Nature Reviews Molecular Cell Biology 25, 187–211. 10.1038/s41580-023-00673-0.

17. Arai, M., Suetaka, S., and Ooka, K. (2024). Dynamics and interactions of intrinsically disordered proteins. Curr Opin Struct Biol 84, 102734. 10.1016/j.sbi.2023.102734.

18. Bondos, S.E., Dunker, A.K., and Uversky, V.N. (2022). Intrinsically disordered proteins play diverse roles in cell signaling. Cell Communication and Signaling 20, 20. 10.1186/s12964-022-00821-7.

19. Majila, K., Ullanat, V., and Viswanath, S. (2026). Disobind: A sequence-based, partner-dependent contact map and interface residue predictor for intrinsically disordered regions. Cell Syst 17, 101486. 10.1016/j.cels.2025.101486.

20. Tabar, M.S., Parsania, C., Giardina, C., Feng, Y., Wong, A.C.H., Metierre, C., Nagarajah, R., Dhungel, B.P., Rasko, J.E.J., and Bailey, C.G. (2025). Intrinsically Disordered Regions Define Unique Protein Interaction Networks in CHD Family Remodelers. FASEB J 39, e70632. 10.1096/fj.202402808RR.

21. Leighton, G.O., Shang, S., Hageman, S., Ginder, G.D., and Williams, D.C., Jr. (2023). Analysis of the complex between MBD2 and the histone deacetylase core of NuRD reveals key interactions critical for gene silencing. Proc Natl Acad Sci U S A 120, e2307287120. 10.1073/pnas.2307287120.

22. Saito, M., and Ishikawa, F. (2002). The mCpG-binding domain of human MBD3 does not bind to mCpG but interacts with NuRD/Mi2 components HDAC1 and MTA2. J Biol Chem 277, 35434–35439. 10.1074/jbc.M203455200.

23. Schmidberger, J.W., Sharifi Tabar, M., Torrado, M., Silva, A.P., Landsberg, M.J., Brillault, L., AlQarni, S., Zeng, Y.C., Parker, B.L., Low, J.K., and Mackay, J.P. (2016). The MTA1 subunit of the nucleosome remodeling and deacetylase complex can recruit two copies of RBBP4/7. Protein Sci 25, 1472–1482. 10.1002/pro.2943.

24. Liu, Z., Ajit, K., Wu, Y., Zhu, W.G., and Gullerova, M. (2024). The GATAD2B-NuRD complex drives DNA:RNA hybrid-dependent chromatin boundary formation upon DNA damage. EMBO J 43, 2453–2485. 10.1038/s44318-024-00111-7.

25. Wunderlich, T.M., Deshpande, C., Paasche, L.W., Friedrich, T., Diegmuller, F., Haddad, E., Kreienbaum, C., Naseer, H., Stebel, S.E., Daus, N., et al. (2024). ZNF512B binds RBBP4 via a variant NuRD interaction motif and aggregates chromatin in a NuRD complex-independent manner. Nucleic Acids Res 52, 12831–12849. 10.1093/nar/gkae926.

26. Reid, X.J., Low, J.K.K., and Mackay, J.P. (2023). A NuRD for all seasons. Trends Biochem Sci 48, 11–25. 10.1016/j.tibs.2022.06.002.

27. Sharifi Tabar, M., Giardina, C., Feng, Y., Francis, H., Moghaddas Sani, H., Low, J.K.K., Mackay, J.P., Bailey, C.G., and Rasko, J.E.J. (2022). Unique protein interaction networks define the chromatin remodelling module of the NuRD complex. Febs j 289, 199–214. 10.1111/febs.16112.

28. Majila, K., Arvindekar, S., Jindal, M., and Viswanath, S. (2025). Frontiers in integrative structural modeling of macromolecular assemblies. QRB Discov 6, e3. 10.1017/qrd.2024.15.

29. Arvindekar, S., Majila, K., and Viswanath, S. (2025). Recent Methods from Statistical Inference for Integrative Structural Modeling. In Springer Handbook of Chem- and Bioinformatics, J. Leszczynski, ed. (Springer Nature Switzerland), pp. 1075–1103. 10.1007/978-3-031-81728-1_47.

30. Olivet, J., Shewakramani, N.R., Cesare, J., Laval, F., Nde, J., Van de Veire, J., Brammerloo, Y., Brebel, B., Debnath, O., Richardson, A.D., et al. (2026). Dynamic engagement of dual-role regulators by the Sin3 complex. bioRxiv, 2026.2002.2026.708192. 10.64898/2026.02.26.708192.

31. Banks, C.A.S., Zhang, Y., Miah, S., Hao, Y., Adams, M.K., Wen, Z., Thornton, J.L., Florens, L., and Washburn, M.P. (2020). Integrative Modeling of a Sin3/HDAC Complex Sub-structure. Cell Rep 31, 107516. 10.1016/j.celrep.2020.03.080.

32. Liu, X., Zhang, Y., Wen, Z., Hao, Y., Banks, C.A.S., Lange, J.J., Slaughter, B.D., Unruh, J.R., Florens, L., Abmayr, S.M., et al. (2020). Driving integrative structural modeling with serial capture affinity purification. Proc Natl Acad Sci U S A 117, 31861–31870. 10.1073/pnas.2007931117.

33. Liu, X., Zhang, Y., Wen, Z., Hao, Y., Banks, C.A.S., Cesare, J., Bhattacharya, S., Arvindekar, S., Lange, J.J., Xie, Y., et al. (2024). An integrated structural model of the DNA damage-responsive H3K4me3 binding WDR76:SPIN1 complex with the nucleosome. Proc Natl Acad Sci U S A 121, e2318601121. 10.1073/pnas.2318601121.

34. Saltzberg, D.J., Viswanath, S., Echeverria, I., Chemmama, I.E., Webb, B., and Sali, A. (2021). Using Integrative Modeling Platform to compute, validate, and archive a model of a protein complex structure. Protein Sci 30, 250–261. 10.1002/pro.3995.

35. Arvindekar, S., Jackman, M.J., Low, J.K.K., Landsberg, M.J., Mackay, J.P., and Viswanath, S. (2022). Molecular architecture of nucleosome remodeling and deacetylase sub-complexes by integrative structure determination. Protein Sci 31, e4387. 10.1002/pro.4387.

36. Low, J.K.K., Silva, A.P.G., Sharifi Tabar, M., Torrado, M., Webb, S.R., Parker, B.L., Sana, M., Smits, C., Schmidberger, J.W., Brillault, L., et al. (2020). The Nucleosome Remodeling and Deacetylase Complex Has an Asymmetric, Dynamic, and Modular Architecture. Cell Rep 33, 108450. 10.1016/j.celrep.2020.108450.

37. Adams, M.K., Banks, C.A.S., Thornton, J.L., Kempf, C.G., Zhang, Y., Miah, S., Hao, Y., Sardiu, M.E., Killer, M., Hattem, G.L., et al. (2020). Differential Complex Formation via Paralogs in the Human Sin3 Protein Interaction Network. Mol Cell Proteomics 19, 1468–1484. 10.1074/mcp.RA120.002078.

38. Gomez, A.V., Galleguillos, D., Maass, J.C., Battaglioli, E., Kukuljan, M., and Andres, M.E. (2008). CoREST represses the heat shock response mediated by HSF1. Mol Cell 31, 222–231. 10.1016/j.molcel.2008.06.015.

39. Kalin, J.H., Wu, M., Gomez, A.V., Song, Y., Das, J., Hayward, D., Adejola, N., Wu, M., Panova, I., Chung, H.J., et al. (2018). Targeting the CoREST complex with dual histone deacetylase and demethylase inhibitors. Nat Commun 9, 53. 10.1038/s41467-017-02242-4.

40. Jumper, J., Evans, R., Pritzel, A., Green, T., Figurnov, M., Ronneberger, O., Tunyasuvunakool, K., Bates, R., Zidek, A., Potapenko, A., et al. (2021). Highly accurate protein structure prediction with AlphaFold. Nature 596, 583–589. 10.1038/s41586-021-03819-2.

41. Honorato, R.V., Trellet, M.E., Jimenez-Garcia, B., Schaarschmidt, J.J., Giulini, M., Reys, V., Koukos, P.I., Rodrigues, J., Karaca, E., van Zundert, G.C.P., et al. (2024). The HADDOCK2.4 web server for integrative modeling of biomolecular complexes. Nat Protoc 19, 3219–3241. 10.1038/s41596-024-01011-0.

42. Del Toro, N., Shrivastava, A., Ragueneau, E., Meldal, B., Combe, C., Barrera, E., Perfetto, L., How, K., Ratan, P., Shirodkar, G., et al. (2022). The IntAct database: efficient access to fine-grained molecular interaction data. Nucleic Acids Res 50, D648–D653. 10.1093/nar/gkab1006.

43. Steinkamp, R., Tsitsiridis, G., Brauner, B., Montrone, C., Fobo, G., Frishman, G., Avram, S., Oprea, T.I., and Ruepp, A. (2025). CORUM in 2024: protein complexes as drug targets. Nucleic Acids Res 53, D651–D657. 10.1093/nar/gkae1033.

44. Balu, S., Huget, S., Medina Reyes, J.J., Ragueneau, E., Panneerselvam, K., Fischer, S.N., Claussen, E.R., Kourtis, S., Combe, C.W., Meldal, B.H.M., et al. (2025). Complex portal 2025: predicted human complexes and enhanced visualisation tools for the comparison of orthologous and paralogous complexes. Nucleic Acids Res 53, D644–D650. 10.1093/nar/gkae1085.

45. Combe, C.W., Graham, M., Kolbowski, L., Fischer, L., and Rappsilber, J. (2024). xiVIEW: Visualisation of Crosslinking Mass Spectrometry Data. J Mol Biol 436, 168656. 10.1016/j.jmb.2024.168656.

46. Vilhais-Neto, G.C., Fournier, M., Plassat, J.L., Sardiu, M.E., Saraf, A., Garnier, J.M., Maruhashi, M., Florens, L., Washburn, M.P., and Pourquie, O. (2017). The WHHERE coactivator complex is required for retinoic acid-dependent regulation of embryonic symmetry. Nat Commun 8, 728. 10.1038/s41467-017-00593-6.

47. Song, Y., Dagil, L., Fairall, L., Robertson, N., Wu, M., Ragan, T.J., Savva, C.G., Saleh, A., Morone, N., Kunze, M.B.A., et al. (2020). Mechanism of Crosstalk between the LSD1 Demethylase and HDAC1 Deacetylase in the CoREST Complex. Cell Rep 30, 2699–2711 e2698. 10.1016/j.celrep.2020.01.091.

48. Zhang, Y., Ng, H.H., Erdjument-Bromage, H., Tempst, P., Bird, A., and Reinberg, D. (1999). Analysis of the NuRD subunits reveals a histone deacetylase core complex and a connection with DNA methylation. Genes Dev 13, 1924–1935. 10.1101/gad.13.15.1924.

49. Millard, C.J., Varma, N., Saleh, A., Morris, K., Watson, P.J., Bottrill, A.R., Fairall, L., Smith, C.J., and Schwabe, J.W. (2016). The structure of the core NuRD repression complex provides insights into its interaction with chromatin. Elife 5, e13941. 10.7554/eLife.13941.

50. Alqarni, S.S., Murthy, A., Zhang, W., Przewloka, M.R., Silva, A.P., Watson, A.A., Lejon, S., Pei, X.Y., Smits, A.H., Kloet, S.L., et al. (2014). Insight into the architecture of the NuRD complex: structure of the RbAp48-MTA1 subcomplex. J Biol Chem 289, 21844–21855. 10.1074/jbc.M114.558940.

51. Cramer, J.M., Scarsdale, J.N., Walavalkar, N.M., Buchwald, W.A., Ginder, G.D., and Williams, D.C., Jr. (2014). Probing the dynamic distribution of bound states for methylcytosine-binding domains on DNA. J Biol Chem 289, 1294–1302. 10.1074/jbc.M113.512236.

52. Farnung, L., Ochmann, M., and Cramer, P. (2020). Nucleosome-CHD4 chromatin remodeler structure maps human disease mutations. Elife 9. 10.7554/eLife.56178.

53. Gnanapragasam, M.N., Scarsdale, J.N., Amaya, M.L., Webb, H.D., Desai, M.A., Walavalkar, N.M., Wang, S.Z., Zu Zhu, S., Ginder, G.D., and Williams, D.C., Jr. (2011). p66Alpha-MBD2 coiled-coil interaction and recruitment of Mi-2 are critical for globin gene silencing by the MBD2-NuRD complex. Proc Natl Acad Sci U S A 108, 7487–7492. 10.1073/pnas.1015341108.

54. Davenport, M.L., Davis, M.R., Davenport, B.N., Crossman, D.K., Hall, A., Pike, J., Harada, S., Hurst, D.R., and Edmonds, M.D. (2022). Suppression of SIN3A by miR-183 Promotes Breast Cancer Metastasis. Mol Cancer Res 20, 883–894. 10.1158/1541-7786.Mcr-21-0508.

55. Lewis, M.J., Liu, J., Libby, E.F., Lee, M., Crawford, N.P., and Hurst, D.R. (2016). SIN3A and SIN3B differentially regulate breast cancer metastasis. Oncotarget 7, 78713–78725. 10.18632/oncotarget.12805.

56. Ismail, H., Chagraoui, J., and Sauvageau, G. (2025). CoREST in pieces: Dismantling the CoREST complex for cancer therapy and beyond. Sci Adv 11, eads6556. 10.1126/sciadv.ads6556.

57. Barrios, A.P., Gomez, A.V., Saez, J.E., Ciossani, G., Toffolo, E., Battaglioli, E., Mattevi, A., and Andres, M.E. (2014). Differential properties of transcriptional complexes formed by the CoREST family. Mol Cell Biol 34, 2760–2770. 10.1128/mcb.00083-14.

58. Burg, J.M., Makhoul, A.T., Pemble, C.W.t., Link, J.E., Heller, F.J., and McCafferty, D.G. (2015). A rationally-designed chimeric KDM1A/KDM1B histone demethylase tower domain deletion mutant retaining enzymatic activity. FEBS Lett 589, 2340–2346. 10.1016/j.febslet.2015.07.028.

59. Da, G., Lenkart, J., Zhao, K., Shiekhattar, R., Cairns, B.R., and Marmorstein, R. (2006). Structure and function of the SWIRM domain, a conserved protein module found in chromatin regulatory complexes. Proc Natl Acad Sci U S A 103, 2057–2062. 10.1073/pnas.0510949103.

60. Kim, S.A., Zhu, J., Yennawar, N., Eek, P., and Tan, S. (2020). Crystal Structure of the LSD1/CoREST Histone Demethylase Bound to Its Nucleosome Substrate. Mol Cell 78, 903–914 e904. 10.1016/j.molcel.2020.04.019.

61. Nde, J., Kempf, C.G., Zimmermann, R.C., Cesare, J., Zhang, Y., Workman, J.L., Florens, L., and Washburn, M.P. (2025). Integrative Structural Modeling of Intrinsically Disordered Regions in a Human HDAC2 Chromatin Remodeling Complex. bioRxiv, 2025.2008.2008.669391. 10.1101/2025.08.08.669391.

62. Kim, D.H., and Han, K.H. (2021). Target-binding behavior of IDPs via pre-structured motifs. Prog Mol Biol Transl Sci 183, 187–247. 10.1016/bs.pmbts.2021.07.031.

63. Lee, S.H., Kim, D.H., Han, J.J., Cha, E.J., Lim, J.E., Cho, Y.J., Lee, C., and Han, K.H. (2012). Understanding pre-structured motifs (PreSMos) in intrinsically unfolded proteins. Curr Protein Pept Sci 13, 34–54. 10.2174/138920312799277974.

64. Russel, D., Lasker, K., Webb, B., Velazquez-Muriel, J., Tjioe, E., Schneidman-Duhovny, D., Peterson, B., and Sali, A. (2012). Putting the pieces together: integrative modeling platform software for structure determination of macromolecular assemblies. PLoS Biol 10, e1001244. 10.1371/journal.pbio.1001244.

65. Grzenda, A., Lomberk, G., Zhang, J.S., and Urrutia, R. (2009). Sin3: master scaffold and transcriptional corepressor. Biochim Biophys Acta 1789, 443–450. 10.1016/j.bbagrm.2009.05.007.

66. Marcum, R.D., and Radhakrishnan, I. (2019). Inositol phosphates and core subunits of the Sin3L/Rpd3L histone deacetylase (HDAC) complex up-regulate deacetylase activity. J Biol Chem 294, 13928–13938. 10.1074/jbc.RA119.009780.

67. Konwar, C., Maini, J., Kohli, S., Brahmachari, V., and Saluja, D. (2022). SIN-3 functions through multi-protein interaction to regulate apoptosis, autophagy, and longevity in Caenorhabditis elegans. Scientific Reports 12, 10560. 10.1038/s41598-022-13864-0.

68. Barnes, V.L., Laity, K.A., Pilecki, M., and Pile, L.A. (2018). Systematic Analysis of SIN3 Histone Modifying Complex Components During Development. Sci Rep 8, 17048. 10.1038/s41598-018-35093-0.

69. Hoffmann, A., and Spengler, D. (2019). Chromatin Remodeling Complex NuRD in Neurodevelopment and Neurodevelopmental Disorders. Frontiers in Genetics 10. 10.3389/fgene.2019.00682.

70. Helness, A., Fraszczak, J., Joly-Beauparlant, C., Bagci, H., Trahan, C., Arman, K., Shooshtarizadeh, P., Chen, R., Ayoub, M., Côté, J.-F., et al. (2021). GFI1 tethers the NuRD complex to open and transcriptionally active chromatin in myeloid progenitors. Communications Biology 4, 1356. 10.1038/s42003-021-02889-2.

71. Lu, Y.A., Sun, J., Wang, L., Wang, M., Wu, Y., Getachew, A., Matthews, R.C., Li, H., Peng, W.G., Zhang, J., et al. (2024). ELM2-SANT Domain-Containing Scaffolding Protein 1 Regulates Differentiation and Maturation of Cardiomyocytes Derived From Human-Induced Pluripotent Stem Cells. J Am Heart Assoc 13, e034816. 10.1161/jaha.124.034816.

72. Abad, C., Robayo, M.C., Muñiz-Moreno, M.d.M., Bernardi, M.T., Otero, M.G., Kosanovic, C., Griswold, A.J., Pierson, T.M., Walz, K., and Young, J.I. (2024). Gatad2b, associated with the neurodevelopmental syndrome GAND, plays a critical role in neurodevelopment and cortical patterning. Translational Psychiatry 14, 33. 10.1038/s41398-023-02678-x.

73. Desai, M.A., Webb, H.D., Sinanan, L.M., Scarsdale, J.N., Walavalkar, N.M., Ginder, G.D., and Williams, D.C., Jr. (2015). An intrinsically disordered region of methyl-CpG binding domain protein 2 (MBD2) recruits the histone deacetylase core of the NuRD complex. Nucleic Acids Res 43, 3100–3113. 10.1093/nar/gkv168.

74. Luo, Y., and Li, H. (2020). Structure-Based Inhibitor Discovery of Class I Histone Deacetylases (HDACs). Int J Mol Sci 21. 10.3390/ijms21228828.

75. Liu, F., Lossl, P., Scheltema, R., Viner, R., and Heck, A.J.R. (2017). Optimized fragmentation schemes and data analysis strategies for proteome-wide cross-link identification. Nat Commun 8, 15473. 10.1038/ncomms15473.

76. Connelly, J.J., Yuan, P., Hsu, H.C., Li, Z., Xu, R.M., and Sternglanz, R. (2006). Structure and function of the Saccharomyces cerevisiae Sir3 BAH domain. Mol Cell Biol 26, 3256–3265. 10.1128/mcb.26.8.3256-3265.2006.

77. He, Y., Imhoff, R., Sahu, A., and Radhakrishnan, I. (2009). Solution structure of a novel zinc finger motif in the SAP30 polypeptide of the Sin3 corepressor complex and its potential role in nucleic acid recognition. Nucleic Acids Res 37, 2142–2152. 10.1093/nar/gkp051.

78. Clark, M.D., Marcum, R., Graveline, R., Chan, C.W., Xie, T., Chen, Z., Ding, Y., Zhang, Y., Mondragon, A., David, G., and Radhakrishnan, I. (2015). Structural insights into the assembly of the histone deacetylase-associated Sin3L/Rpd3L corepressor complex. Proc Natl Acad Sci U S A 112, E3669–3678. 10.1073/pnas.1504021112.

79. Xie, T., He, Y., Korkeamaki, H., Zhang, Y., Imhoff, R., Lohi, O., and Radhakrishnan, I. (2011). Structure of the 30-kDa Sin3-associated protein (SAP30) in complex with the mammalian Sin3A corepressor and its role in nucleic acid binding. J Biol Chem 286, 27814–27824. 10.1074/jbc.M111.252494.

80. Forneris, F., Binda, C., Adamo, A., Battaglioli, E., and Mattevi, A. (2007). Structural basis of LSD1-CoREST selectivity in histone H3 recognition. J Biol Chem 282, 20070–20074. 10.1074/jbc.C700100200.

81. Tochio, N., Umehara, T., Koshiba, S., Inoue, M., Yabuki, T., Aoki, M., Seki, E., Watanabe, S., Tomo, Y., Hanada, M., et al. (2006). Solution structure of the SWIRM domain of human histone demethylase LSD1. Structure 14, 457–468. 10.1016/j.str.2005.12.004.

82. Viswanath, S., and Sali, A. (2019). Optimizing model representation for integrative structure determination of macromolecular assemblies. Proc Natl Acad Sci U S A 116, 540–545. 10.1073/pnas.1814649116.

83. Bonomi, M., Hanot, S., Greenberg, C.H., Sali, A., Nilges, M., Vendruscolo, M., and Pellarin, R. (2019). Bayesian Weighing of Electron Cryo-Microscopy Data for Integrative Structural Modeling. Structure 27, 175–188 e176. 10.1016/j.str.2018.09.011.

84. Ullanat, V., Kasukurthi, N., and Viswanath, S. (2022). PrISM: precision for integrative structural models. Bioinformatics 38, 3837–3839. 10.1093/bioinformatics/btac400.

85. Pettersen, E.F., Goddard, T.D., Huang, C.C., Meng, E.C., Couch, G.S., Croll, T.I., Morris, J.H., and Ferrin, T.E. (2021). UCSF ChimeraX: Structure visualization for researchers, educators, and developers. Protein Sci 30, 70–82. 10.1002/pro.3943.

86. van Zundert, G.C.P., Rodrigues, J., Trellet, M., Schmitz, C., Kastritis, P.L., Karaca, E., Melquiond, A.S.J., van Dijk, M., de Vries, S.J., and Bonvin, A. (2016). The HADDOCK2.2 Web Server: User-Friendly Integrative Modeling of Biomolecular Complexes. J Mol Biol 428, 720–725. 10.1016/j.jmb.2015.09.014.

87. Humphrey, W., Dalke, A., and Schulten, K. (1996). VMD: Visual molecular dynamics. Journal of Molecular Graphics 14, 33–38. 10.1016/0263-7855(96)00018-5.

