## Supplemental Information for "Multicomplex Integrative Structural Modeling of a Human Histone Deacetylase Interactome"

**Description of Supplementary Tables**

**Supplementary Table S1. Cross linking mass spectrometry results of HDAC1-Halo and HDAC2-Halo Affinity Purifications. A.** Crosslink Spectrum Matches (CSMs) from DSSO-treated affinity-purifications of HEK293T cells stably expressing HDAC1-Halo. **B.** Crosslink Spectrum Matches (CSMs) from DSSO-treated affinity-purifications of HEK293T cells stably expressing HDAC2-Halo. **C.** Non-Redundant Crosslinked Pairs Detected in HDAC1-Halo and/or HDAC2-Halo XLMS Data. **D.** Non-Redundant Proteins Detected in HDAC1-Halo and/or HDAC2-Halo XLMS Data **E.** Input Values for Figures 1 A-C and S1C.

**Supplementary Table S2. Detailed list of all interaction interfaces within each of the protein complexes as modeled by the Integrative Modeling Platform.**

**Supplementary Table S3. NuRD subcomplex modeling data.** Crosslinking data used from both HDAC1 and HDAC2 analyses used to build Figure 7 and Supplemental Figure 12.

**Supplementary Table S4. Comparison of secondary structures in the NuRD subcomplex across 3D models.** Details of secondary structures from AlphaFold compared to the integrated structural model for HDAC1, MBD3, MTA1, GATAD2B, and RBBP4.

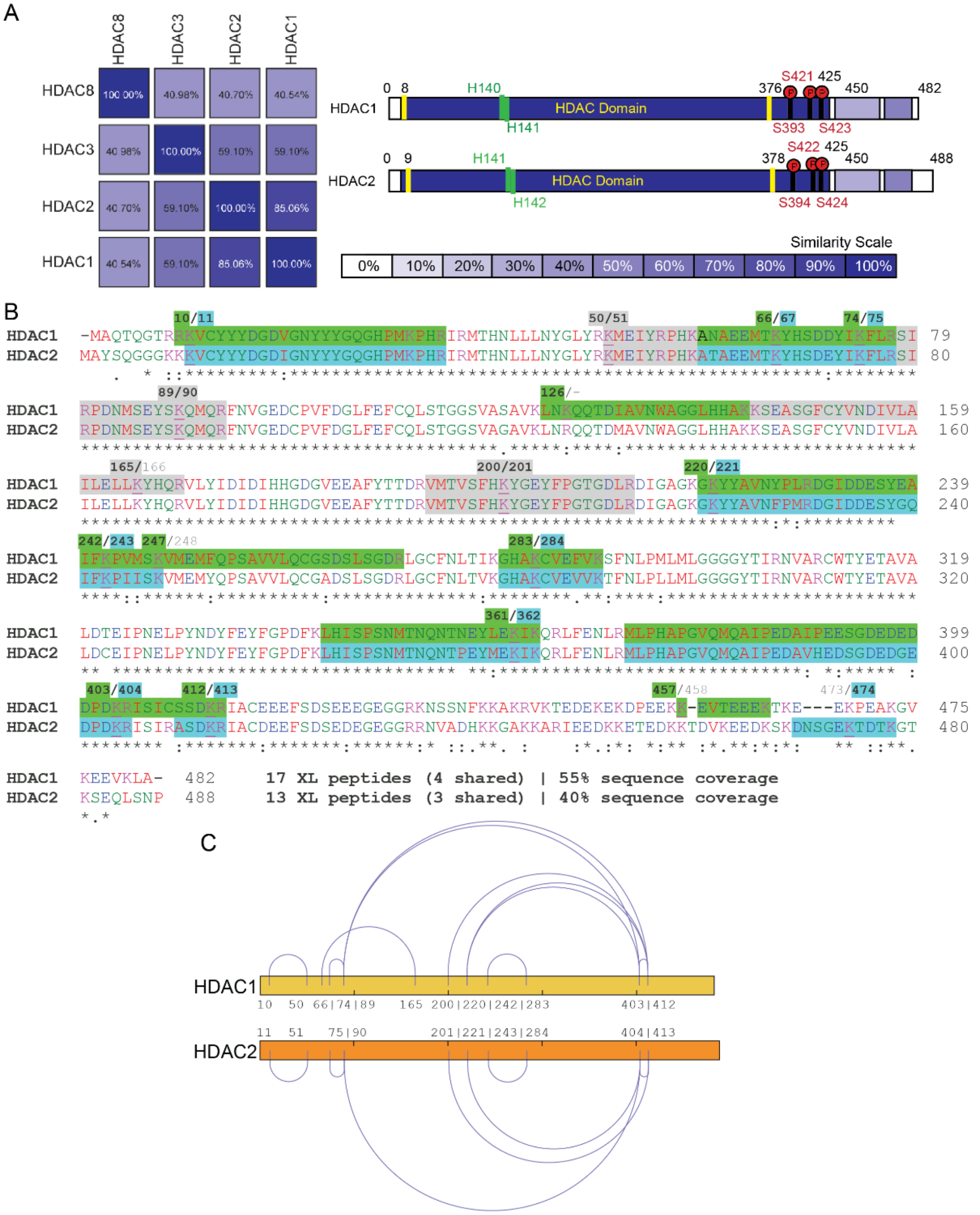


**Figure S1: Sequence homology between HDAC1 and HDAC2 and detected crosslinked peptides**. **A.** Class I HDAC sequence similarities and percent overlaps at the amino acid level between HDAC1 and HDAC2 as completed and visualized by Lalnprot at Expasy. The known HDAC domain (yellow), Zn2+ active sites (green), and phosphorylation sites (red) are highlighted. **B.** Alignment of HDAC1 and HDAC2 sequences with CLUSTAL Omega (1.2.4), such as the symbols below the sequences indicate exact matches (*), strongly (:) and weakly (.) similar substitutions. The detected crosslinked peptides are mapped in green and blue for peptides unique to HDAC1 and HDC2, respectively, while shared peptides are shown in gray. Crosslinked lysine residues are underlined and numbered above the sequences (see Table S1A-B). **C.** Intra-molecular crosslinks within HDAC1 and 2 visualized by xiView (see Table S1E).

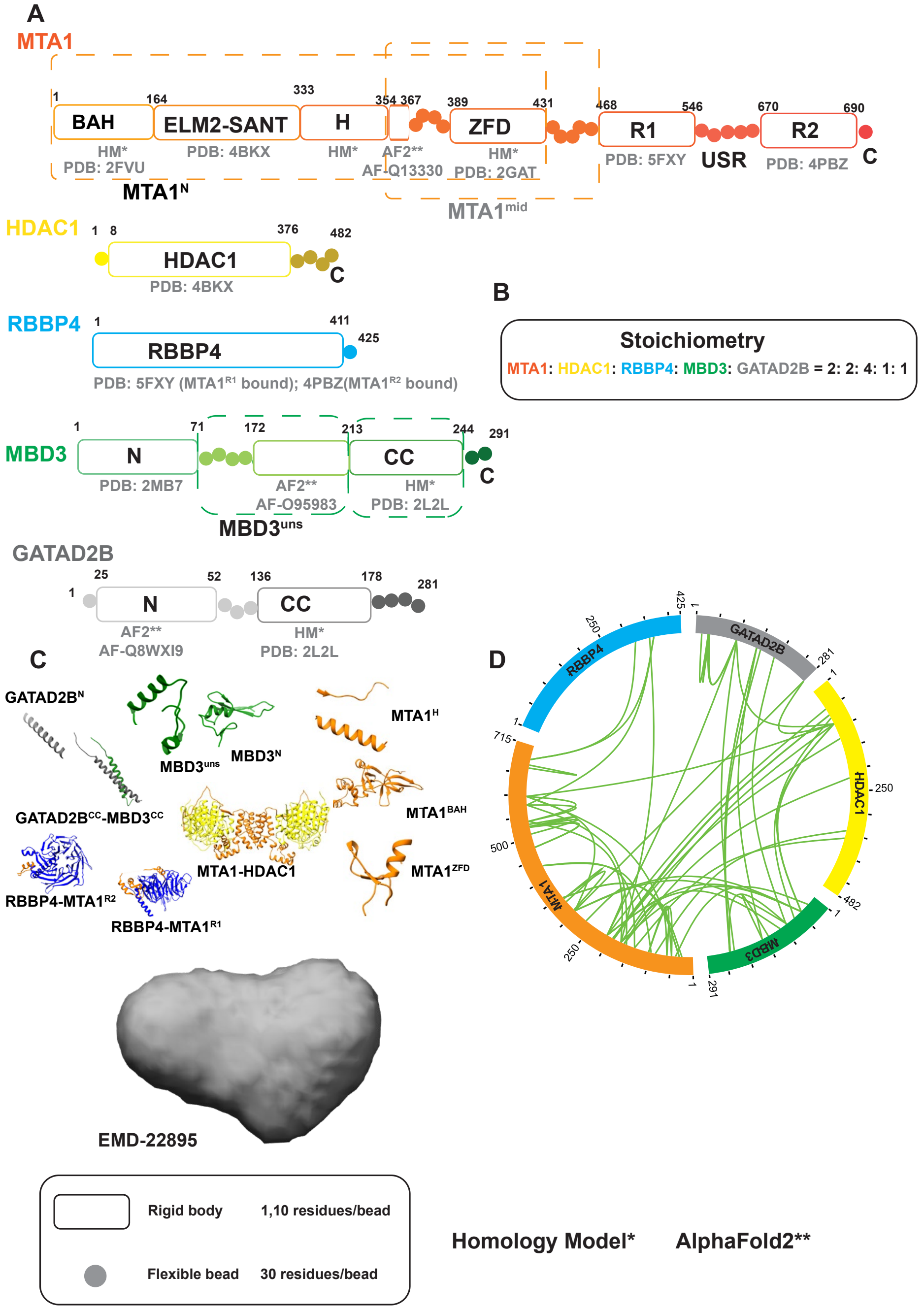


**Figure S2: Structural information gathered to model the NuRD complex with IMP. A.** Domains of the isoforms modeled in the study are shown in progressively darker shades along their sequence. Regions for which atomic structures exist, or can be predicted, are represented by rectangles, whereas regions without known structure are represented by beads. PDB IDs are shown for existing subunit structures and templates of homology models (HM); similarly, Alphafold Database IDs are shown where AF2 was used. Dashed rectangles represent domain names. **B.** Atomic structures and EM maps used for modeling. PDB and Alphafold IDs are given in A. **C.** Crosslinks used for integrative modeling, represented as an xiView circos plot. **D.** Stoichiometry of the modeled 5-subunit sub-complex.

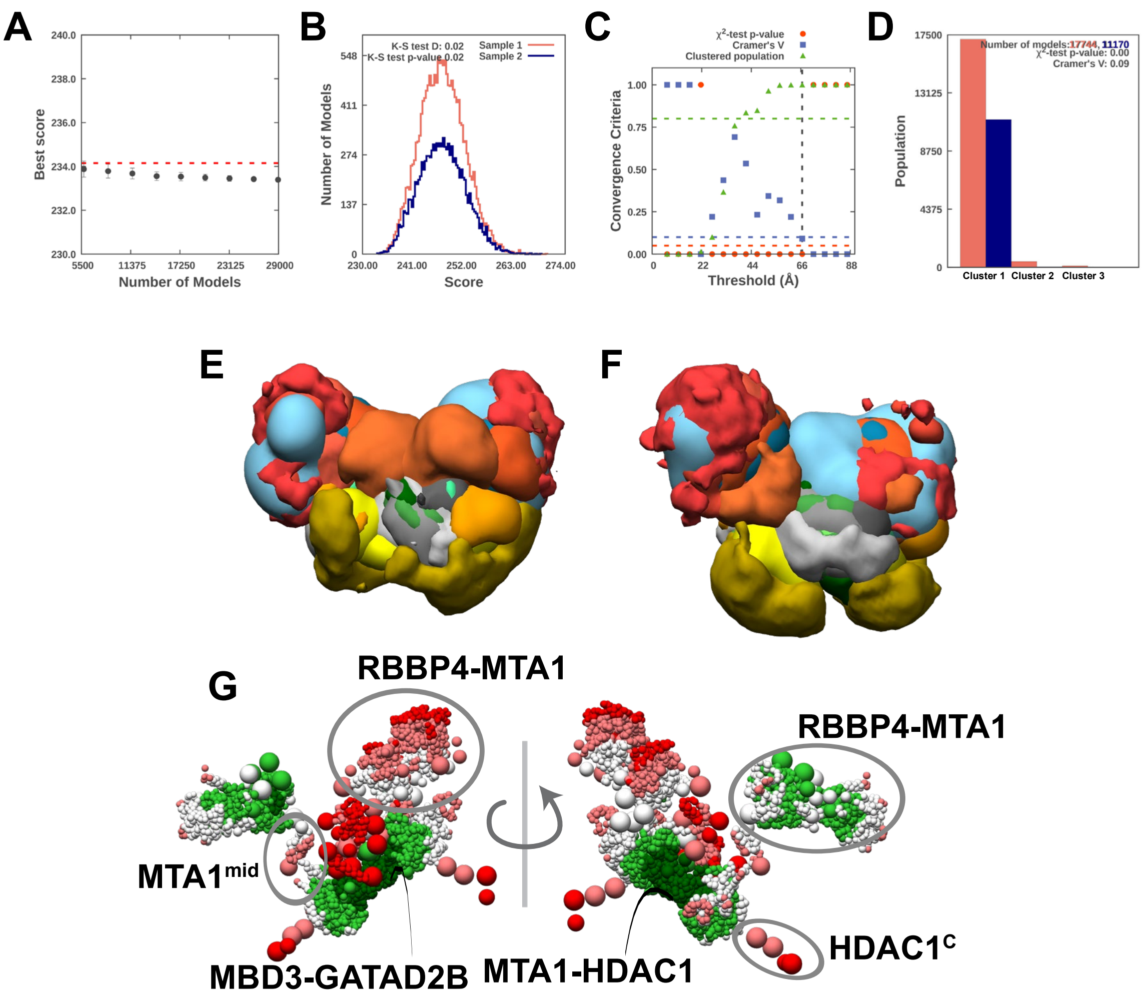


**Figure S3. Sampling exhaustiveness protocol and precision analysis for NuRD integrative models. A.** The graph highlights the convergence of the model score for the 28,914 good-scoring models. The scores do not continue to improve as more models are computed essentially independently. The error bars represent the standard deviations of the best scores estimated by iterating sampling of models for 10 cycles. The red dotted line indicates a lower bound reference on the total score. **B.** The distribution graph shows the similarity of model score between sample 1 (red) and 2 (blue). The difference in the distribution of scores is significant (Kolmogorov-Smirnov two-sample test p-value is <0.05), however the magnitude of the difference is small (Kolmogorov-Smirnov two-sample test statistic D is <0.03). Hence, the two score distributions are effectively equal. **C.** The plot shows three criteria for determining the sampling precision (Y-axis) evaluated as a function of the RMSD clustering threshold (X-axis). The considered criteria are: (i) the p-value, computed using the $X^2$-test for homogeneity of proportions (red dots); (ii) an effect size for the $X^2$-test, quantified by the Cramer's V value (blue squares); and (iii) the population of models in sufficiently large clusters (containing at least 10 models from each sample) as shown as green triangles. The vertical dotted grey line indicates the RMSD clustering threshold at which all three conditions are satisfied (p-value > 0.05; dotted red line), Cramer's V <0.10 (dotted blue line), and the population of clustered models >0.80 (dotted green line), thus defining the sampling precision as 67 Å. **D.** The bar graph depicts the population of models in sample 1 and 2 for the clusters obtained by threshold-based clustering (RMSD threshold=67 Å). **E-F.** The comparison of localization probability densities for NuRD from samples 1 and 2 in the major cluster are shown. The cross-correlation of the density maps for the two samples is greater than 0.95. **G.** Precision of integrative models was analyzed using PrISM. Regions of high precision are represented in green while regions of low precision are represented in red.

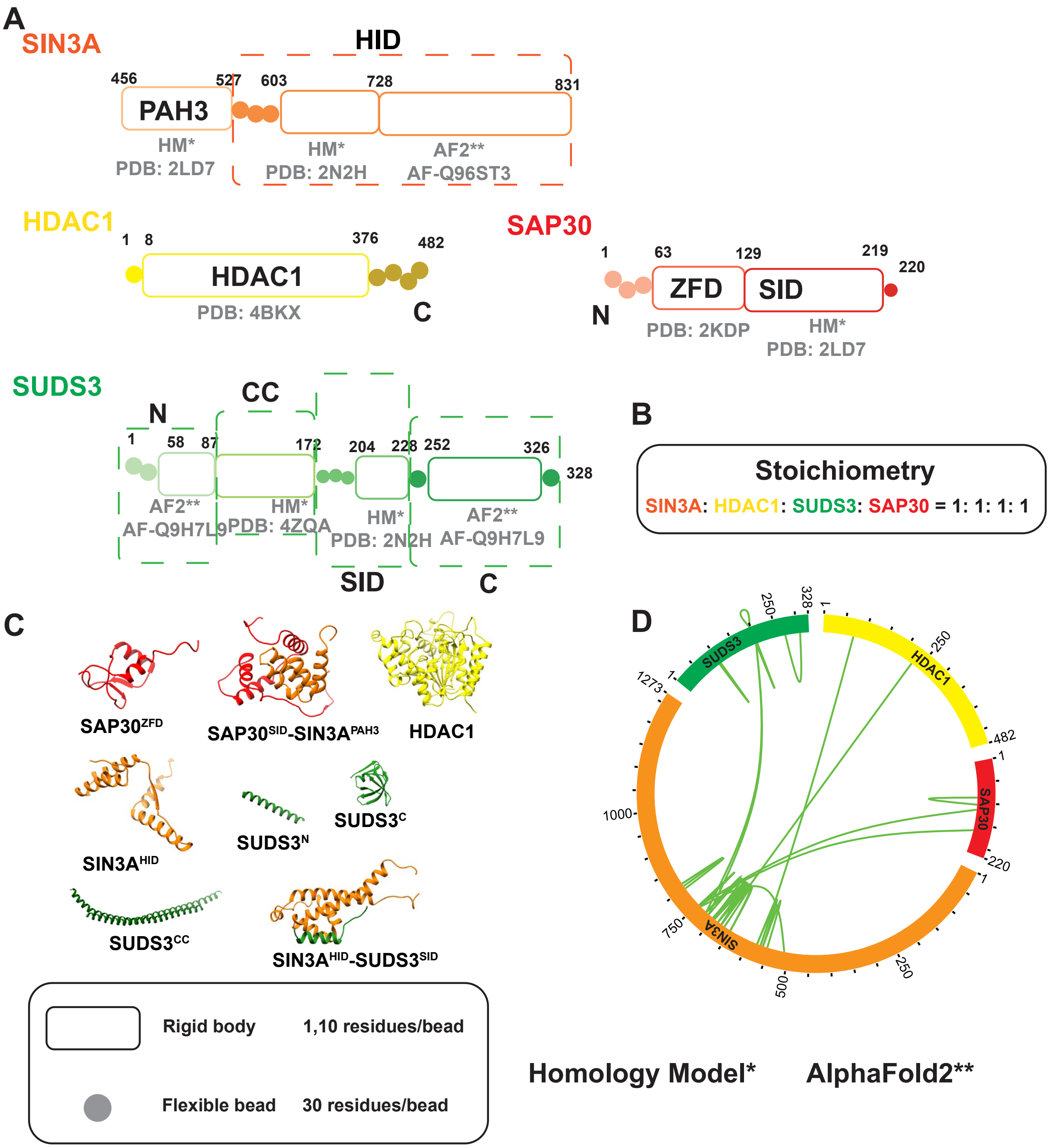


**Figure S4. Structural information gathered to model the SIN3A complex with IMP. A.** Domains of the isoforms modeled in the study are shown in progressively darker shades along their sequence. Regions for which atomic structures exist, or can be predicted, are represented by rectangles, whereas regions without known structure are represented by beads. PDB IDs are shown for existing subunit structures and templates of homology models (HM); similarly, Alphafold Database IDs are shown where AF2 was used. Dashed rectangles represent domain names. **B.** Atomic structures and EM maps used for modeling. PDB and Alphafold IDs are given in A. **C.** Crosslinks used for integrative modeling, represented as an xiView circos plot. **D**. Stoichiometries of the modeled 4-subunit sub-complex.

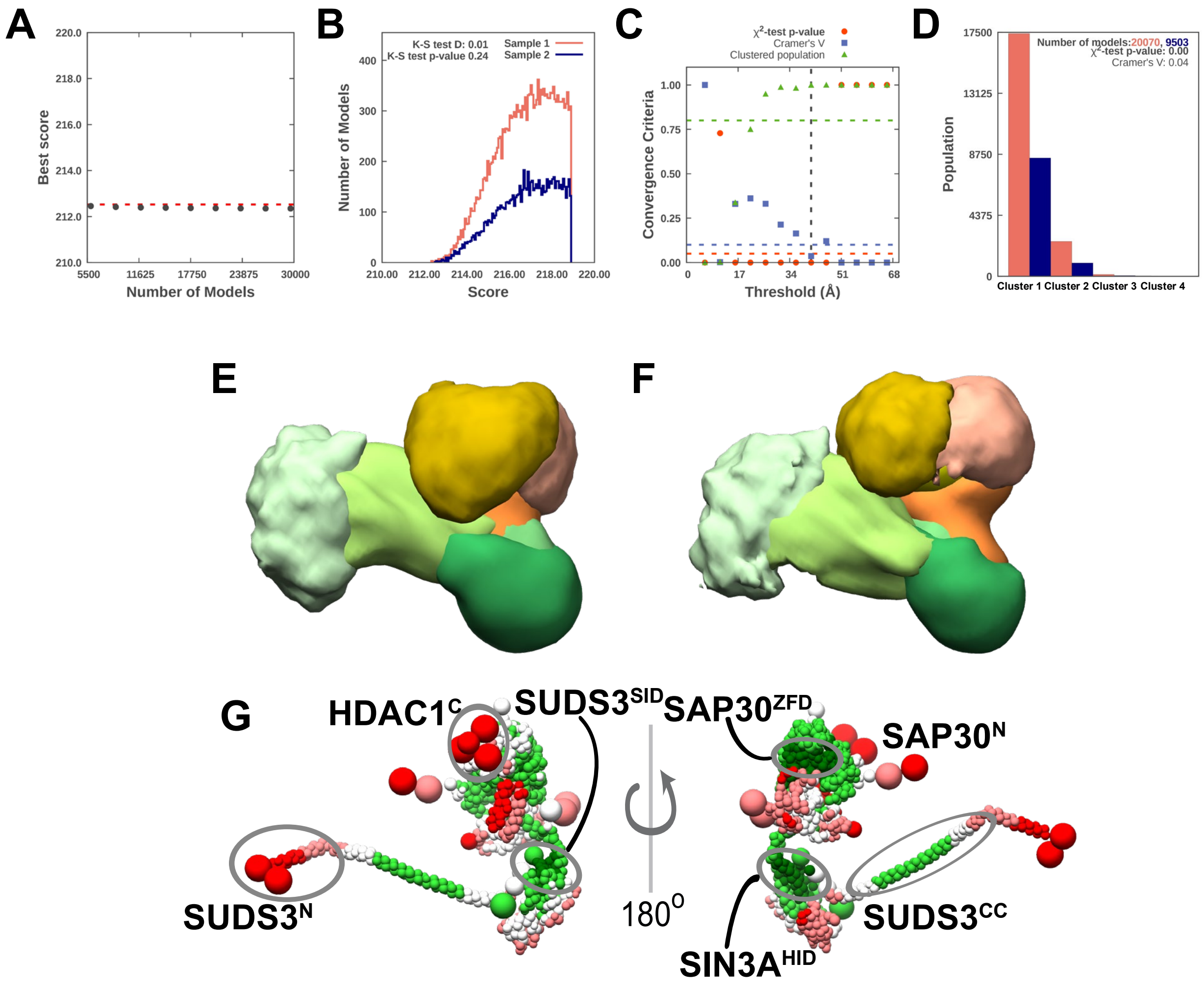


**Figure S5. Sampling exhaustiveness protocol and precision analysis for SIN3A integrative models. A.** The graph highlights the convergence of the model score for the 29,602 good-scoring models. The scores do not continue to improve as more models are computed essentially independently. The error bar represents the standard deviations of the best scores estimated by iterating sampling of models for 10 cycles. The red dotted line indicates a lower bound reference on the total score. **B.** The distribution graph shows the testing similarity of model score between sample 1 (red) and 2 (blue). The difference in the distribution of scores is not significant (Kolmogorov-Smirnov two-sample test p-value is <0.05), and the magnitude of the difference is small (Kolmogorov-Smirnov two-sample test statistic D is <0.03). Hence, the two score distributions are effectively equal. **C.** The plot shows three criteria for determining the sampling precision (Y-axis) evaluated as a function of the RMSD clustering threshold (X-axis). The considered criteria are: (i) the p-value, computed using the $X^2$-test for homogeneity of proportions (red dots); (ii) an effect size for the $X^2$-test, is quantified by the Cramer's V value (blue squares); and (iii) the population of models in sufficiently large clusters (containing at least 10 models from each sample), shown as green triangles. The vertical dotted grey line indicates the RMSD clustering threshold at which all three conditions are satisfied (p-value > 0.05; dotted red line), Cramer's V <0.10 (dotted blue line), and the population of clustered models >0.80 (dotted green line), thus defining the sampling precision as 41 Å. **D.** The bar graph depicts the population of models in sample 1 and 2 for the clusters obtained by threshold-based clustering (RMSD threshold=41 Å). **E-F.** The comparison of localization probability densities for SIN3A from samples 1 & 2 in the major cluster are shown. The cross-correlation of the density maps for the two samples is greater than 0.95. **G.** Precision of integrative models was analyzed using PrISM. Regions of high precision are represented in green while regions of low precision are represented in red.

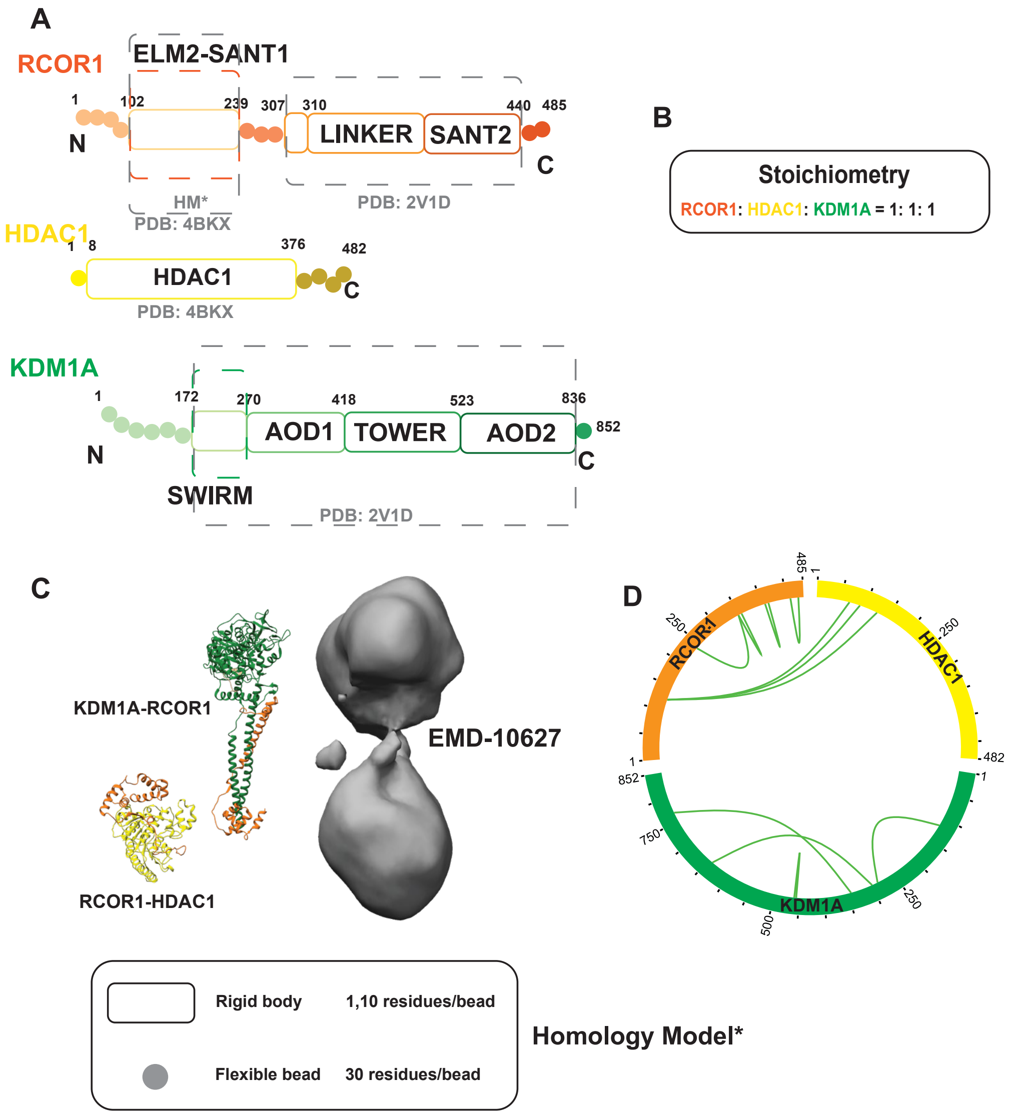


**Figure S6. Structural information gathered to model the CoREST complex with IMP. A.** Domains of the isoforms modeled in the study are shown in progressively darker shades along their sequence. Regions for which atomic structures exist, or can be predicted, are represented by rectangles, whereas regions without known structure are represented by beads. PDB IDs are shown for existing subunit structures and templates of homology models (HM); similarly Alphafold Database IDs are shown where AF2 was used. Dashed rectangles represent domain names. **B.** Atomic structures and EM maps used for modeling. PDB and Alphafold IDs are given in A. **C.** Crosslinks used for integrative modeling, represented as an xiView circos plot. **D.** Stoichiometries of the modeled 5-subunit sub-complex.

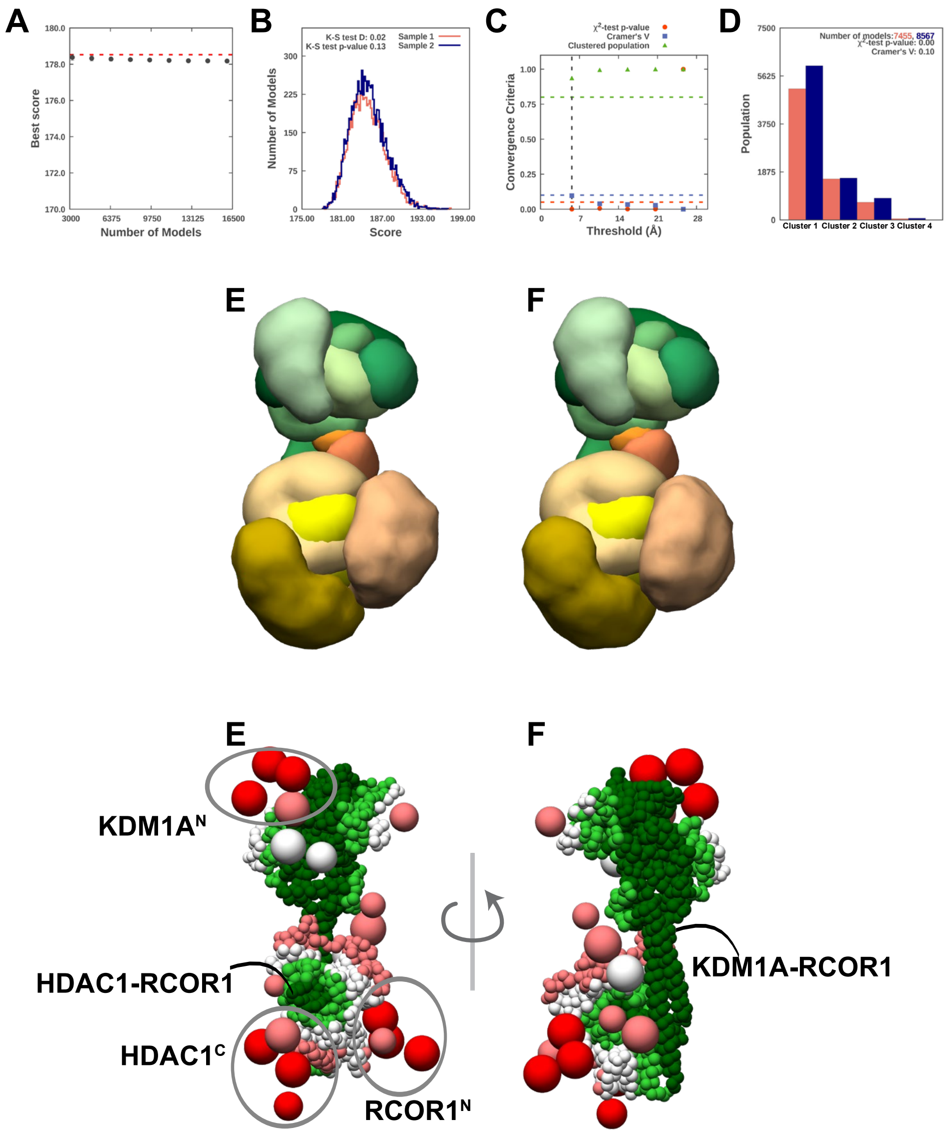


**Figure S7. Sampling exhaustiveness protocol and precision analysis of CoREST integrative models. A.** The graph highlights the convergence of the model score for the 16,055 good-scoring models. The scores do not continue to improve as more models are computed essentially independently. The error bar represents the standard deviations of the best scores estimated by iterating sampling of models for 10 cycles. The red dotted line indicates a lower bound reference on the total score. **B.** The distribution graph shows the testing similarity of model score between sample 1 (red) and 2 (blue). The difference in the distribution of scores is not significant (Kolmogorov-Smirnov two-sample test p-value is <0.05), and the magnitude of the difference is small (Kolmogorov-Smirnov two-sample test statistic D is <0.03). Hence, the two score distributions are effectively equal. **C.** The plot shows three criteria for determining the sampling precision (Y-axis) evaluated as a function of the RMSD clustering threshold (X-axis). The considered criteria are: (i) the p-value, computed using the $X^2$-test for homogeneity of proportions (red dots); (ii) an effect size for the $X^2$-test, quantified by the Cramer's V value (blue squares); and (iii) the population of models in sufficiently large clusters (containing at least 10 models from each sample), shown as green triangles. The vertical dotted grey line indicates the RMSD clustering threshold at which all three conditions are satisfied (p-value > 0.05; dotted red line), Cramer's V <0.10 (dotted blue line), and the population of clustered models >0.80 (dotted green line), thus defining the sampling precision as 5 Å. **D.** The bar graph depicts the population of models in sample 1 and 2 for the clusters obtained by threshold-based clustering (RMSD threshold=15 Å). **E-F.** The comparison of localization probability densities for CoREST from sample A & B in the major cluster are shown. The cross-correlation of the density maps for the two samples is greater than 0.95. **G.** Precision of integrative models was analyzed using PrISM. Regions of high precision are represented in green while regions of low precision are represented in red.

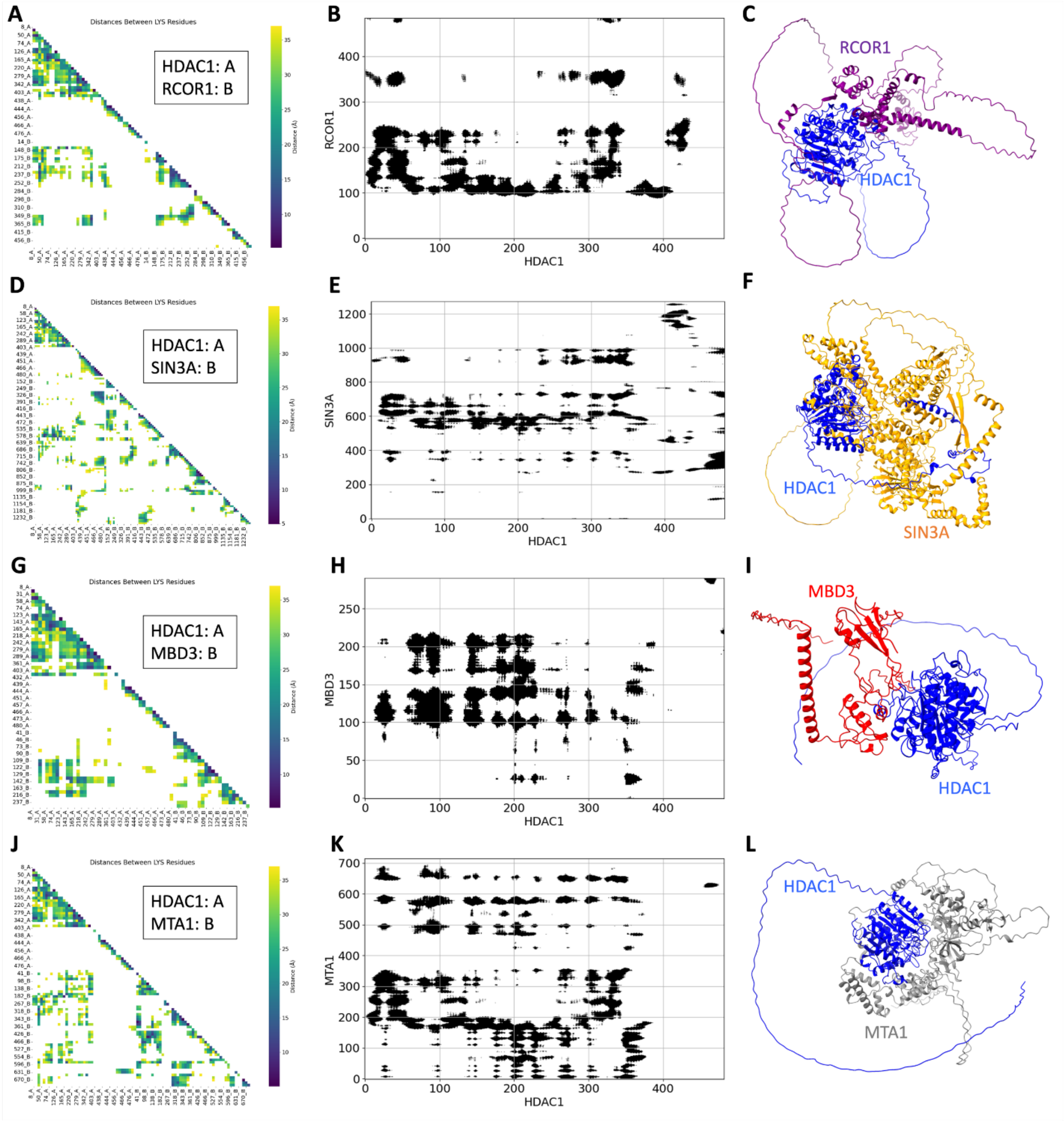


**Figure S8: Interaction and structural analysis of AlphaFold-only dimeric complexes**. (A-C) *HDAC1/RCOR1 complex; (D-F) HDAC1/SIN3A complex; (G-I) HDAC1/MBD3 complex; (J-L) HDAC1/MTA1 complex.* (A, D, G, and J) Lysine-lysine interaction maps for different complexes, highlighting key hotspots of the lysine-lysine interactions. (B, E, H, and K) show the residue contact maps distribution of interacting residues in each dimer. (C, F, I, and L) represent the 3D visualization of the different dimers, allowing for comparison of the overall structural conformation. These individual structures of the dimeric complexes highlight their unique structural features and the unique HDAC1 conformations in the N- and C-termini.

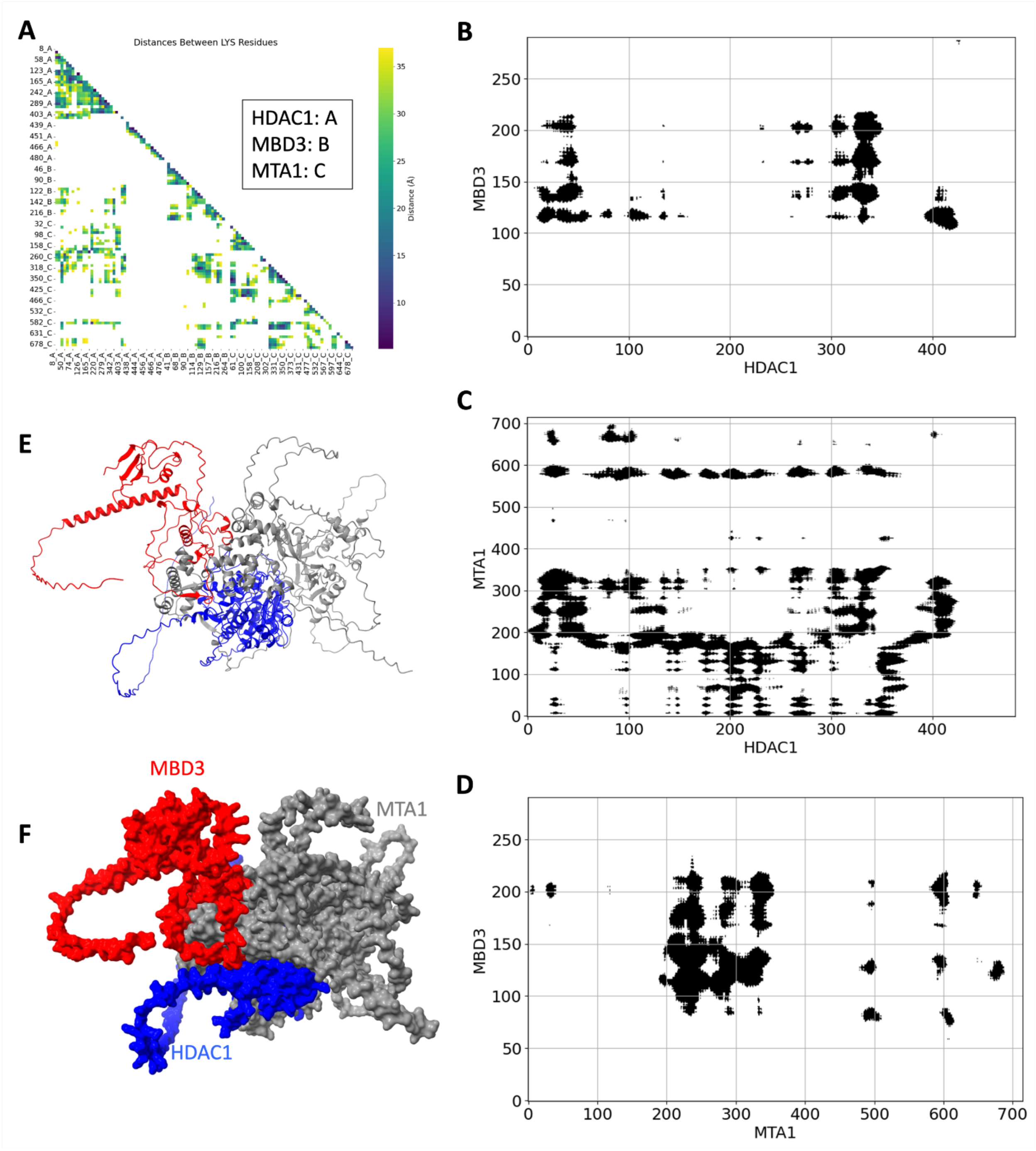


**Figure S9**: **Structural and interaction analysis of the AlphaFold-only trimeric HDAC1/MBD3/MTA1 complex**. (A) Lysine-lysine interaction maps for the HDAC1/MBD3/MTA1 trimer, illustrating key lysine residues between all three components. (B-D) Residue contact maps showing the interactions between HDAC1 and MBD3 (B), HDAC1 and MTA1 (C), or MTA1 and MBD3 (D) within the trimeric complex. (E, F) 3D visualization of the HDAC1/MBD3/MTA1 trimer and its surface representation (F) showcasing the unique conformation of the HDAC1 in the context of the trimeric complex.

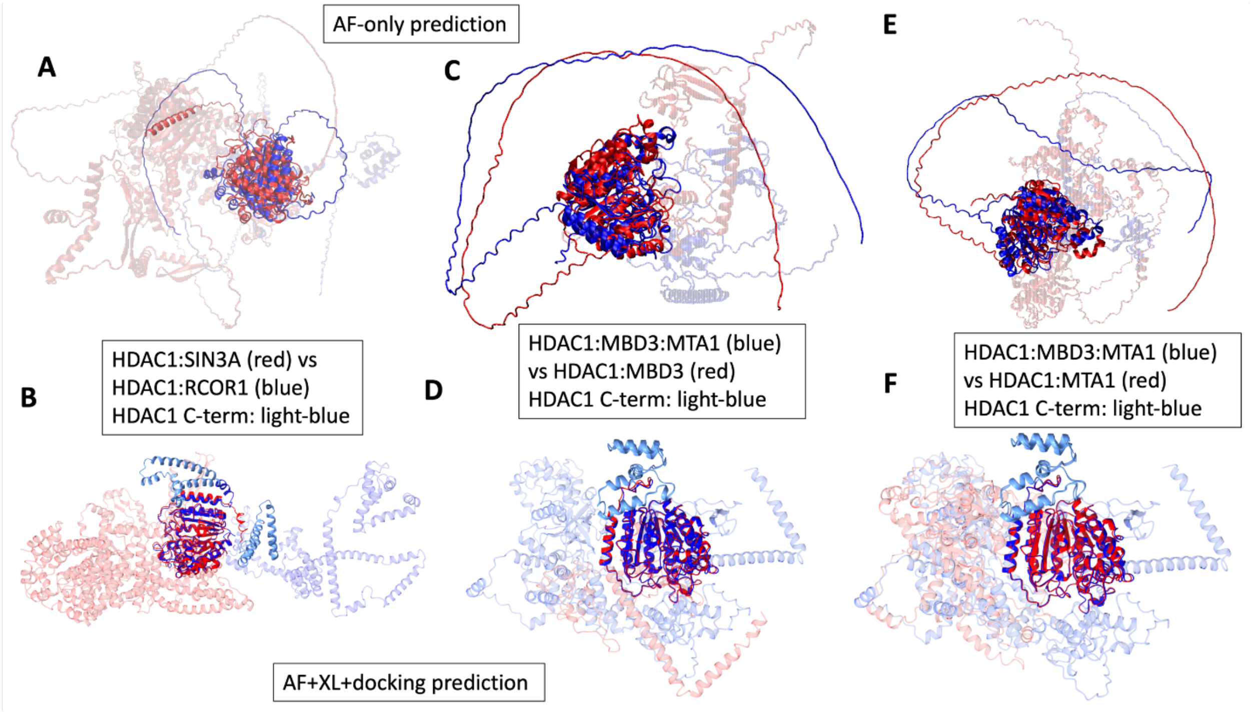


**Figure S10**. **Comparison of HDAC1 structures in dimers and trimers from both approaches providing insights into flexibility and complex formation**. (A, B) Superimposed structure of HDAC1 from the dimeric HDAC1/RCOR1 (blue) and HDAC1/SIN3A (red) complexes determined by AlphaFold-only (A) and integrative (B). It highlights the greater flexibility in the C-terminal region of HDAC1 (shown in light blue) when bound to RCOR1 compared to SIN3A. (C, D) Superimposed structure of HDAC1 from the trimeric HDAC1/MBD3/MTA1 complex (blue) and the dimeric HDAC1/MBD3 complex (red) determined by AlphaFold-only (C) and integrative (D). It demonstrates a more compact conformation in the trimer, as compared to the more flexible dimer (C) but with less flexible C-terminal of HDAC1 (shown in light blue) in both cases (D). (E, F) Superimposed structure of HDAC1 from the trimeric HDAC1/MBD3/MTA1 complex (blue) and the dimeric HDAC1/MTA1 complex (red), determined by AlphaFold-only (E) and integrative (F). The trimer aligns better with cross-linking mass spectrometry (XL-MS) observations, supporting the importance of predicting the structure of HDAC1 in the context of larger, multi-protein complexes. These comparisons emphasize not only HDAC1 folds differently (specifically in the C-terminal region shown in light blue) in the presence of each protein but also the need to consider interactions with more than two additional proteins for an accurate structural model of HDAC1.

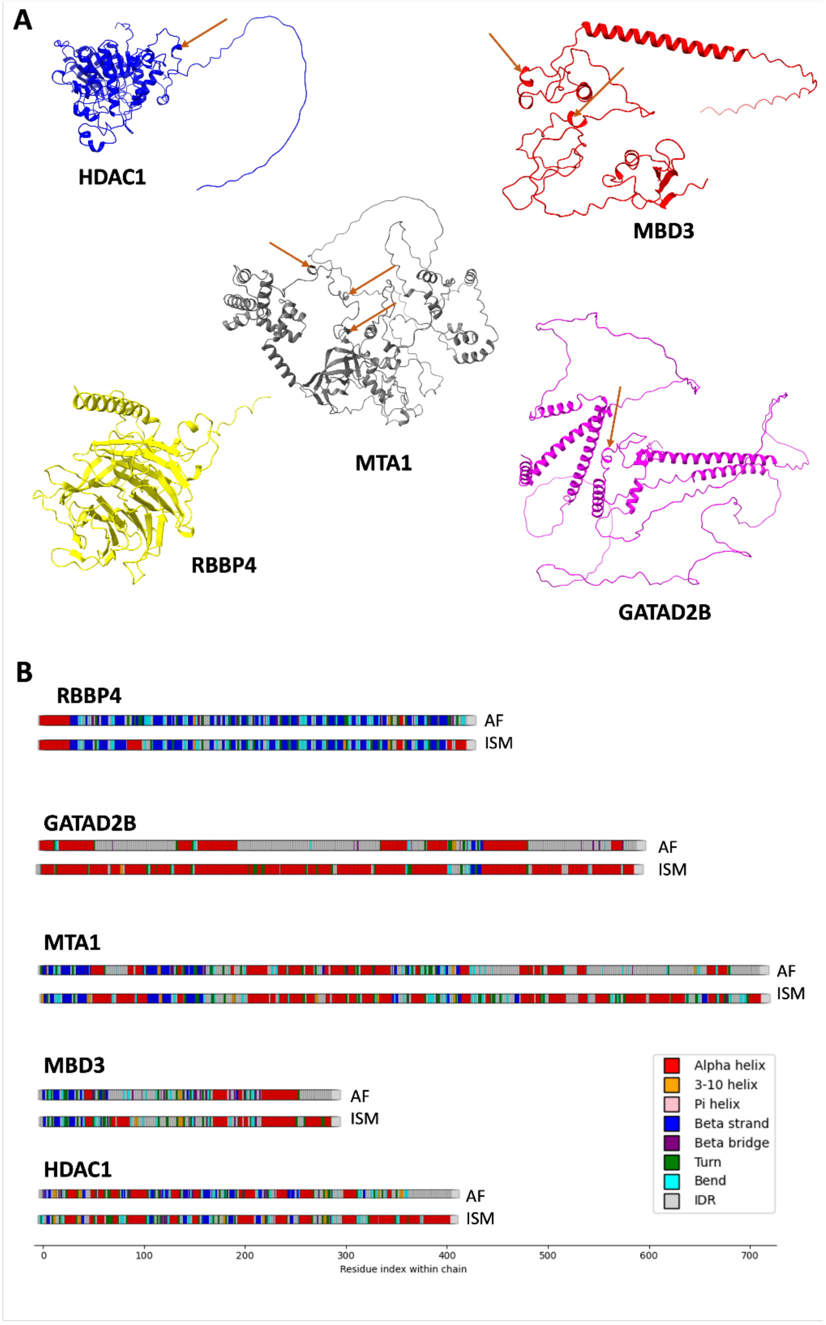


**Figure S11. Structural analysis of individual NuRD complex components in monomeric and complexed forms.** (A) Predicted secondary structures of the five NuRD complex proteins—HDAC1 (chain A), MBD3 (chain B), MTA1 (chain C), GATAD2B (chain D), and RBBP4 (chain E)—in their monomeric forms, highlighting the presence of IDRs. (B) Secondary structures of the same proteins within the assembled complex, showing a notable increase in α-helical content, indicating that several IDRs adopt ordered conformations upon complex formation. (C) 3D representations of the individual proteins, with arrows marking pre-structured motifs within the IDRs, suggesting regions predisposed to fold upon interaction.

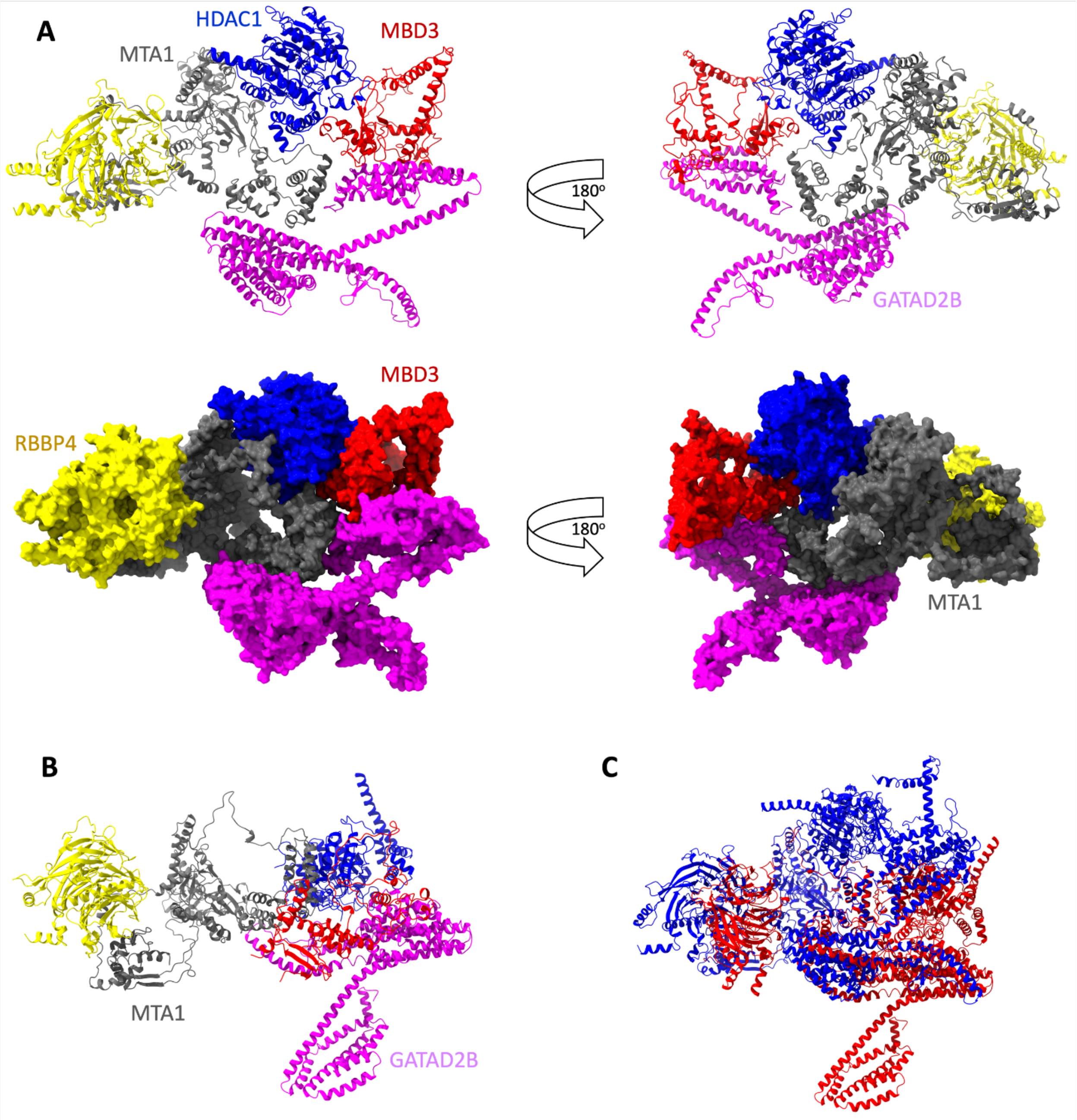


**Figure S12. 3D Representation of the HDAC1:MBD3:MTA1:GATAD2B:RBBP4 complex obtained via Integrative Structural Modeling. A.** Initial structural model of the NuRD subcomplex shown in both cartoon and surface representation. This model satisfies 87% of the experimentally observed crosslinks, indicating a high level of agreement with crosslinking mass spectrometry data. **B.** Refined model generated by incorporating the unmatched crosslinks from panel A, providing an alternative conformation of the complex. **C.** Superposition of the initial and refined models, highlighting regions of structural flexibility within the NuRD complex and illustrating the dynamic architecture of the spatial arrangement of its subunits.
